# Chronic SSRI treatment reverses HIV-1 protein-mediated synaptodendritic damage

**DOI:** 10.1101/2021.01.11.426213

**Authors:** Adam R. Denton, Charles F. Mactutus, Almeera U. Lateef, Steven B. Harrod, Rosemarie M. Booze

## Abstract

HIV-1 infection affects approximately 37 million individuals and approximately 50% of seropositive individuals will develop symptoms of clinical depression and apathy. Dysfunctions of both serotonergic and dopaminergic neurotransmission have been implicated in the pathogenesis of motivational alterations. The present study evaluated the efficacy of a SSRI (escitalopram) in the HIV-1 transgenic (Tg) rat. Behavioral, neurochemical, and neuroanatomical outcomes with respect to HIV-1 and sex were evaluated to determine the efficacy of chronic escitalopram treatment. Escitalopram treatment restored function in each of the behavioral tasks that were sensitive to HIV-1 induced impairments. Further, escitalopram treatment restored HIV-1-mediated synaptodendritic damage in the nucleus accumbens; treatment with escitalopram significantly increased dendritic proliferation in HIV-1 Tg rats. However, restoration did not consistently occur with the neurochemical analysis in the HIV-1 rat. Taken together, these results suggest a role for SSRI therapies in repairing long-term HIV-1 protein-mediated neuronal damage and restoring function.

## Introduction

In the United States, approximately 50% of HIV-infected individuals will experience symptoms of clinical depression and/or apathy throughout their lifetime (Savetsky et al., 2001; Rabkin et al., 2008; Campos et al., 2010; Bhatia and Munjal, 2014, Castellon et al., 1998). The incidence of major depressive disorder in HIV seropositive individuals is roughly twice that of HIV seronegative individuals (Do et al., 2014 Pence et al., 2018; Arseniou et al., 2014; Mills et al., 2018). Comorbid depression remains a serious impediment to the successful treatment of HIV (Farinpour et al., 2003), with depression significantly impacting adherence to combination antiretroviral therapy (cART) and medical appointment attendance (Horberg et al., 2008; Pence et al., 2018; Yoo-Jeong et al., 2016).

Apathy remains a frequent psychological disturbance among HIV seropositive individuals, despite cART treatment. The persistence of apathetic symptoms despite treatment is not surprising given the close association between apathetic behavioral responses and clinical depression (Marin, Firinciogullari, and Biedrzycki, 1993), however, lending support to the notion that apathy and depression are dissociable is that they share a differential relationship, a dissociation, from neurocognitive performance in HIV seropositive subjects Castellon, et al., 1998). In addition to the described symptoms, roughly half of all individuals with HIV will develop some HIV-associated neurocognitive disorders (HAND) (Sanmarti, 2014; Bryant et al., 2015).

The development of apathy is a direct effect of HIV infection (McIntosh et al., 2015), proposed as a consequent to transactivator of transcription (Tat) and envelope glycoprotein (gp120) protein exposure (Bertrand et al., 2018). These proteins have been shown to produce harmful effects upon the neural circuitry underlying reward pathways, and the dopaminergic system more specifically (Illenberger et al., 2020). Dopaminergic dysfunction accompanying HIV infection has been examined in human brain tissue (Silvers et al., 2006; Kumar et al., 2011 Purohit et al., 2011), cell culture systems (Aksenov et al., 2008; Bertrand et al., 2013) and in animals used to model symptoms of HIV infection. (Fitting et al., 2015; Javadi-Paydar et al., 2017; Bertrand et al., 2018; Denton et al., 2019). HIV-1 induced dopaminergic disruption may play a critical role in apathy, which has been documented in the HIV-1 Tg rat, in response to DAT dysfunction and decreased dopamine levels (Javadi-Paydar et al.,2017; Bertrand et al., 2018; Denton et al., 2019).

The HIV-1 Tg rat brain contains seven of the nine genes that comprise the HIV viral genome, resulting in a non-infectious, long-term model of HIV-1 viral protein exposure (Reid et al., 2001; Vigorito et al., 2015 McLaurin et al., 2018). The HIV-1 Tg rat was initially generated using an infectious provirus derivation following the deletion of the *Sph1-Bal1* fragment that encompasses the *gag and pol* genes of the virus, rendering the HIV-1 Tg rat non-infectious (Reid et al., 2001). Production of proteins, such as tat and gp120 proteins, remains under the control of the LTR promoter. Viral proteins, such as tat, remain present in cerebrospinal fluid of HIV seropositive individuals despite suppressive antiretroviral therapy and thus continue the cycle of active transcription in the brain, in addition to producing oxidative stress and neuronal injury (Henderson et al., 2019). Moreover, the persistence of HIV-1 infected cells in the brain is associated with decreased neurocognitive performance despite long-term antiretroviral adherence (Spudich et al., 2019). Exposure to viral proteins tat and gp120 is further implicated in synaptic loss, which is present in both clinical populations and animal models (Toggas et al., 1994; Kim et al., 2008; Fitting et al., 2008; Bertrand et al., 2013; Bertrand et al., 2014; Festa et al., 2020).

The HIV-1 Tg rat has previously been demonstrated to have compromised synaptodendritric connectivity in the nucleus accumbens core region (Roscoe et al., 2014; McLaurin et al., 2018), and additionally, compromised dopaminergic and serotonergic function in the nucleus accumbens core and prefrontal cortex, respectively (Denton et al., 2019). Roscoe et al., (2014) reported a profound decrease in dendritic branching complexity in medium spiny neurons of the nucleus accumbens, which is sex-dependent (McLaurin et al., 2018). Moreover, it was reported that HIV-1 Tg rats demonstrated a distributional shift in spine length with HIV-1 Tg animals exhibiting shorter spine lengths in addition to decreased spine volume (Roscoe et al., 2014). These findings were extended by McLaurin et al., (2018), where it was reported that HIV-1 Tg rats exhibited a distributional shift in spine type, with an increased frequency of dendritic spines closer to the soma relative to more distal dendritic branches (McLaurin et al., 2018). HIV-1 induced alterations in synaptic connectivity may occur partially in response to viral proteins and inflammatory cytokines (Kim et al., 2008; Green et al., 2019). Moreover, synaptic loss has been associated with HAND (Ellis et al., 2009).

The impact of these alterations in dendritic complexity in the HIV-1 Tg rat has been demonstrated by *in-vivo* studies of neurotransmission in the HIV-1 Tg rat. Using fast-scan cyclic voltammetry techniques, HIV-1 Tg animals demonstrated diminished peak release of both extracellular dopamine in the nucleus accumbens and serotonin in the prefrontal cortex. Moreover, HIV-1 Tg rats exhibited altered reuptake kinetics for both dopamine and serotonin, relative to control animals (Denton et al., 2019), indicating reductions in DAT and SERT synaptic function. Collectively, these findings illustrate the highly reproducible structural and neurochemical changes that are present within the reward circuitry of the HIV-1 Tg rat (Illenberger et al., 2020), indicating a stable environment for testing neuroprotective and restorative therapeutic approaches.

Escitalopram is a commonly prescribed antidepressant. Escitalopram is a selective serotonin reuptake inhibitor (SSRI), and is one of the most selective SSRIs available, with an approximately 50% greater potency relative to R-citalopram (Braestrup and Sanchez, 2004). Escitalopram acts via the primary serotonin binding site of the serotonin transporter (SERT), in addition to the allosteric regulatory binding site of the SERT. Consequently, escitalopram is an effective medication for serotonin dysregulation (Braestrup and Sanchez, 2004). Moreover, studies examining acute escitalopram treatment in mice have demonstrated efficacy in increasing evoked serotoninergic response. Saylor et al., (2019) examined the effects of acute escitalopram administration using fast-scan cyclic voltammetry (FSCV). Following an acute dosage of escitalopram, a significant (50%) increase in serotonin response was observed (Saylor et al., 2019). In the current study chronic dosing of escitalopram was used, as long-term use of SSRI more fully characterizes typical SSRI treatment in the context of HIV-1 infection.

The present study examined (1) behavioral effects of escitalopram treatment, (2) real-time extracellular release and reuptake kinetics of dopamine and serotonin as measured by FSCV, and finally, (3) morphologic alterations in the nucleus accumbens in the HIV-1 Tg rat. Specifically, the behavioral effects of SSRI treatment in HIV-1 Tg and F344/N rats were determined using a five bottle choice sucrose concentration test, a modified hole board response, an elevated plus-maze task, pre-pulse inhibition (PPI) of the visual PPI and acoustic startle task, and a social behavior task. FSCV was performed following the conclusion of behavioral testing to evaluate dopamine and serotonin kinetics in vivo. Following sacrifice, dendritic complexity and branching of medium spiny neurons (MSN) in the nucleus accumbens were examined using confocal microscopy. Taken together, the present study sought to determine the functional and mechanistic profile of chronic escitalopram treatment in the HIV-1 Tg rat, and establish the potential of SSRI therapeutics in treatment of HIV-1.

## Materials and Methods

### Overall Experimental Design (Figure 1)

Animals were implanted with a subcutaneous pellet of escitalopram and allowed to recover for one week. Following recovery, animals were tested for sucrose preference in the second week. In the third week, exploratory behavior was measured using a modified hole board task. Pre-pulse inhibition testing occurred during the fourth week, followed by elevated plus maze testing and social behavior testing in the fifth week (Denton, 2019). Following the conclusion of behavioral testing, animals were randomly assigned to undergo FSCV. Cage-paired animals not assigned to voltammetry studies were sacrificed for dendritic spine analysis.

**Figure 1:**
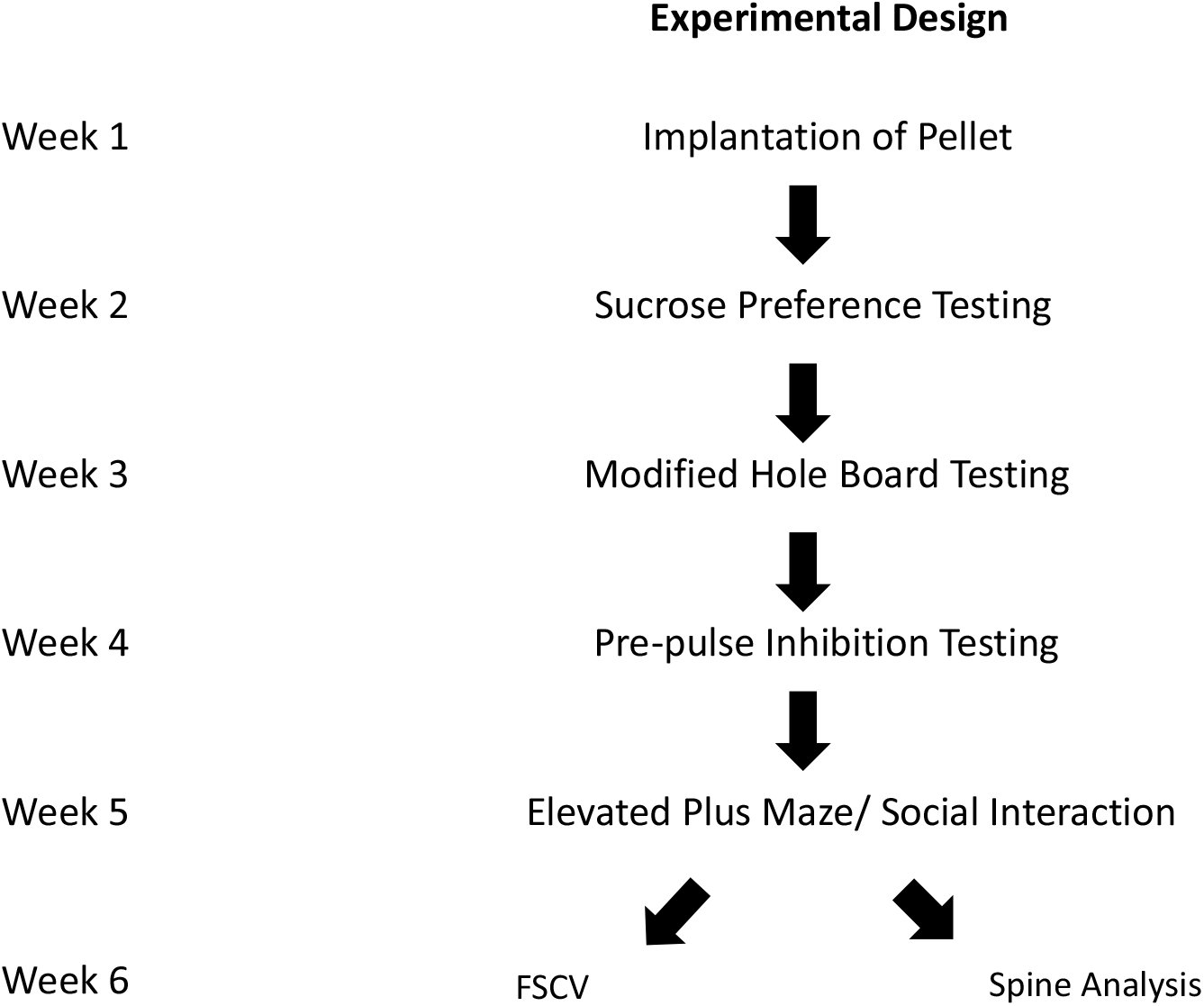
Experimental Design. Rats (N=73; HIV-1 Tg, n=31; F344/N, n=42; Males, n=36; Females, n=44) were implanted with a subcutaneous pellet of escitalopram and allowed to habituate to the 14.76 mg pellet of escitalopram for 1 week. Following this initial week, rats were tested in each behavioral task during the subsequent 4 weeks. Directly after the conclusion of behavioral testing, rats were implanted with electrodes for FSCV recording (n=40) or sacrificed for dendritic spine analysis (n=33).

### Subjects

Adult animals (N=73; HIV-1 Tg, n=31; F344/N, n=42; Males, n=36; Females, n=44) were obtained from Envigo, (Indianapolis, IN) and pair-housed under targeted conditions of 21°± 2° C, 50 % ± 10% relative humidity with a 12 hour light: dark (0700:1900 hours) cycle. Animals were pair-housed by both sex and genotype. Food (Pro-Lab Rat, Mouse, and Hamster chow # 3000) and water were available *ad libitum* throughout the experiment. All behavioral tasks were conducted during diurnal hours and behavioral testing commenced at approximately 6 months of age.

### Drug Treatment

Escitalopram (14.76 mg pellet = 4mg/kg for 40 days) (Sigma Aldrich, Saint Louis, MO) or placebo pellets (Innovative Research of America, Sarasota, FL) were subcutaneously implanted in the medial neck area of each animal. In brief, animals were anesthetized using a 2-3% concentration of Sevoflurane (Henry Schein Animal Health, Dublin, OH). A small (approximately 3 mm) subcutaneous pocket was made into which the pellet was placed. Incisions were then sutured and each animal was administered butorphanol and placed in a recovery chamber with a heating pad. Animals were monitored for one-week post-operatively before beginning behavioral testing.

### Estrous Cycle Tracking

Vaginal lavage was performed at 0900 hours on each day of the testing period to determine the cycle stage of female rodents. Each lavage was performed with approximately 1 mL of freshly prepared phosphate-buffered saline solution. The solution was administered to the vagina of the rat with a standard eyedropper and quickly retracted. The solution was then evaluated under a low-power light microscope to determine the cycle stage via cell type morphology (Booze et al., 1999; Westwood, 2008). All female rodents were behaviorally tested and sacrificed during the morning of diestrus.

### Methods-Behavioral Analyses

#### Sucrose Preference

Animals were individually placed in an empty testing chamber with free access to 0%, 1%, 5%, 10% and 30% concentrations of sucrose solution in 100 ml graduated cylinders equipped with stopper and drinking tube (Ancare, Bellmore, NY). Habituation to the five bottles of sucrose occurred two consecutive days prior to the testing period using distilled H_2_0 in place of sucrose (Denton, 2019). Following habituation, animals were tested in the morning at 1000 hours for 30 minutes per day across five consecutive days. Sucrose consumption was measured both with respect to the meniscus and cylinder weight. Cylinder order was randomized daily using a Latin square design to control for any effect of cylinder position upon sucrose consumption (Bertrand et al., 2018; Denton et al., 2019).

#### Modified Hole Board

A custom made insert equipped with 16 equidistant 3.17 cm holes was placed inside a 40 cm^3^ locomotor activity chamber. Nose pokes into each hole were recorded by photocells placed below the custom insert. Each nose poke was recorded by FlexField Software (San Diego Instruments, San Diego CA). Following a 10 minute habituation period, recording sessions occurred for 10 minutes each day for 7 consecutive days. The apparatus was cleaned with a 10% ethanol solution following each testing period. Testing was performed in the presence of 70db background white noise in a darkened room to encourage exploratory behavior at approximately 1030 hours.

#### Elevated Plus Maze

Each animal received a single testing session in a 109 cm X 109 cm (2 open arm X 2 closed arm) elevated plus-maze apparatus. Behavior was recorded by a camera mounted above the apparatus. Overall activity was recorded with SMART tracking software (San Diego Instruments, San Diego CA). The apparatus was cleaned with a 10% ethanol solution between consecutive trials. Female animals were tested while in diestrus to control for any effect of estrus cycle upon the exploration of the animal. Animals with failed trials were retested one week later. The dependent measure was the time spent in the open arm of the apparatus out of a 10 minute session. Recording sessions occurred at approximately 1100 hours.

#### Visual and Auditory Pre-pulse Inhibition of the Acoustic Startle Response

Animals were placed in a startle chamber (SR-Lab Startle Reflex System, San Diego Instruments) enclosed in an isolation cabinet (Industrial Acoustic Company) and acclimated to the presence of 70dB background noise for 5 minutes at approximately 1400 hours. The subjects were then presented with a series of six pulse-only trials at 100dB (Denton, 2019). Following this acclimation period, subjects were presented with 36 prepulse trials of 85dB with interstimulus intervals (ISIs) of 0, 8, 40, 80, 120, and 4,000 milliseconds assigned in a Latin-square procedure. The stimulus occurred for 20 milliseconds. The 0 and 4000-millisecond intervals were included to provide a baseline acoustic startle response. The apparatus was cleaned thoroughly with a 10% ethanol solution between each session.

#### Social and Play Behavior

On the day of testing, animals were habituated to the testing room for 10 minutes. Animals were then placed into an empty testing chamber at approximately 1300 hours with a bodyweight and sex-matched novel partner. Rodent interaction was recorded for 10 minutes. Total interaction time was recorded as a dependent measure of social behavior. Females were tested in diestrus. Successive trials were conducted in previously cleaned cages to account for any bias due to novel olfactory cues or debris.

### Methods-Fast-Scan Cyclic Voltammetry

#### Manufacture of Carbon Fiber Microelectrodes

Carbon fiber microelectrodes were manufactured by aspirating 7 μm diameter carbon-fibers (Goodfellow Inc, Coraopolis, PA) into glass capillaries (0.6 mm external diameter, 0.4 mm internal diameter, A-M Systems Inc., Sequim, WA). Fibers were sealed into the capillaries with a vertical pipette puller (Narishige Group, Tokyo, Japan). The exposed fiber was trimmed to approximately 50 μm for evaluation of dopamine and precisely 150 μm under a low-light power microscope for evaluation of serotonin, as this length is critical for proper measurement of serotonin (Hashemi et al. 2009; Denton et al., 2019). Nafion, a cation exchange polymer, was electrodeposited onto the carbon fiber portion of each serotonin electrode and dried for 10 minutes at 70° C (Hashemi et al. 2009; Denton et al., 2019).

#### Implantation of Carbon Fiber Microelectrodes and Stimulating Pin

Animals (n=40) were deeply anesthetized using 2-4% Sevoflurane approximately 6 weeks after being implanted with either escitalopram or placebo pellets. The animal’s head was placed into a stereotaxic apparatus (David Kopf Instruments, Tujunga, CA.), with a heating pad to maintain constant body temperature (Denton, 2019). Carbon fiber microelectrodes were placed into both the nucleus accumbens (AP: +2.6, ML: +1.6, DV: −5.8) and CA1/CA2 region of the hippocampus (AP: −5.5, ML: +5.0, DV: −4.0), for evaluation of dopamine and serotonin, respectively (Paxinos and Watson, 2014). A stainless steel stimulating electrode (Plastics One, Roanoke VA) was implanted in the medial forebrain bundle (AP: −2.8, ML: +1.7, DV: −8.0), while a silver reference electrode was placed in the hemisphere contralateral to the stimulating electrode. To stimulate the release of dopamine and serotonin, biphasic pulse trains were applied through a stimulus isolator (NL800A, Neurolog; Medical Systems Corp., Great Neck, NY). To evaluate the release of dopamine, a triangular waveform ranging from −0.4 volts to 1.3 volts in amplitude was applied, while a triangular waveform ranging from 0.2 volts to 1.0 volts in amplitude was used for the evaluation of serotonin. Background-subtracted cyclic voltammograms were obtained as time vs. voltage (x-axis by y-axis). For both neurotransmitters, stimulation parameters were held at a frequency of 60 Hz, with 120 total stim pulses spaced 4ms apart and a stim to scan delay of 89.50ms. Following the conclusion of the recording session, animals were sacrificed under deep anesthesia. Brains were extracted and sectioned to verify current electrode placement.

### Methods-Ballistic Labeling of Medium Spiny Neurons in the Nucleus Accumbens

#### Preparation of Tezfel Tubing

Tezful tubing (IDEX Health Sciences, Oak Harbor, WA) was cut and cleaned with a solution of polyvinylpyrrolidone (PVP) (EMD Millipore Corporation, Billerica, MA) and distilled H_2_0 and allowed to sit at room temperature before use.

#### Preparation of DiOlistic Cartridges

Cartridges were constructed as previously described (Roscoe et al., 2014). Briefly, tungsten beads (Bio-Rad, Hercules, CA) and crystallized Dil (Invitrogen, Carlsbad, CA) were dissolved in methylene chloride (Sigma-Aldrich, St. Louis, MO). Tungsten bead solution was applied to a standard glass slide before being treated with Dil solution and mixed until air-dried. The mixture was then removed from the slides and combined with distilled H_2_0 prior to probe sonication with a Branson Sonifier 150 (Branson Ultrasonics, Danbury, CT). The solution was then drawn into the previously prepared Tezfel tubing and placed into a tubing prep station (Bio-Rad, Hercules, CA) for rotation until even distribution of the tungsten was achieved. The remaining liquid was drawn from the tubing with a syringe and nitrogen gas was blown through the tubing to ensure drying. The tubing was then cut into 13 mm segments and stored in a light-proof container.

#### Ballistic Labeling of Medium Spiny Neurons

Animals (N=33) that underwent behavioral testing (and not used for voltammetry) were sacrificed *via* transcardial perfusion (Variable speed peristaltic pump number 70730-064, VWR, Avantor) of approximately 100 mL of freshly prepared paraformaldehyde approximately 6 weeks after having been implanted with escitalopram or placebo pellets. Brains were then removed and stored in paraformaldehyde. All terminal sacrifices of female rats were conducted during the diestrus phase of the rat estrous cycle. Brains were sliced on a standard rat brain matrix (Ted Pella, Inc., Redding, CA) at a thickness of 500 μm.

Five slices were taken from the nucleus accumbens of each animal and labeled with the Helios Gene Gun (Bio-Rad, Hercules, CA). Previously prepared cartridges were delivered at 70 psi through 3 um pore filter papers onto the tissue. Prepared slices were then washed with PBS and allowed to incubate at 4°C overnight. The following morning, all tissue was mounted and cover-slipped with Fisherbrand 22X50-1.5 glass coverslips (Fisher Scientific, Pittsburgh PA)(Roscoe et al., 2014; McLaurin et al., 2018). Slices were imaged with a Nikon TE-2000E confocal microscope (pinhole size 30 μm, pinhole projected radius 167 nm) using a green helium-neon laser with an emission of 533 nm (Nikon, Tokyo, Japan). Three neurons were imaged at both 20x and 60x magnification. 60x (n.a. = 1.4) images were traced for dendritic and spine complexity using NeuroLucida 360 (MBF Biosciences, Williston, TX). One neuron per animal was used to evaluate spine parameters using Neurolucida Explorer (MBF Biosciences, Williston, TX). Dendritic spines were classified according to backbone length using an algorithm internal to Neurolucida 360 (Rodriguez et al., 2008). Length (μm), volume (μm^3^), and head diameter (μm) were evaluated for each neuron. Spine lengths were defined as between .01 μm and 4 μm (Blanpied and Ehlers, 2004; Ruszczycki et al., 2012) while spine volume was measured between 0.02 μm^3^ and 0.2 μm^3^ (Merino-Serrais et al., 2013; McLaurin et al., 2018). Spine head diameter was defined as between 0.3 μm and 1.2 μm (Bae et al., 2012). A Sholl analysis was performed to examine dendritic complexity as measured by the number of intersections at successive 10 μm radii (Sholl, 1953).

### Data Analysis

Statistical analyses were performed using analysis of variance (ANOVA) and regression techniques (SPSS Statistics 25, IBM Corp., Somer, NY; BMDP statistical software (release 8.1, Statistical Solutions Ltd, Cork, Ireland; SAS/STAT Software 9.4, SAS Institute, Inc., Cary, NC; GraphPad Software, Inc., Version 5.02, La Jolla, CA), where the alpha criterion of *p*≤0.05 was considered to be statistically significant. Orthogonal decompositions the Greenhouse-Geisser df correction factor and/or logarithmic transformations were utilized to address potential violations of the compound symmetry assumption. Based on the *a priori* aims of the present study, planned comparisons were conducted to evaluate the impact of chronic HIV-1 viral protein exposure (i.e., F344/N placebo vs. HIV-1 Tg placebo), the effect of escitalopram treatment in restoring function (i.e., HIV-1 Tg escitalopram vs. HIV-1 Tg placebo), and the magnitude of the escitalopram effect (i.e., HIV-1 escitalopram vs. F344/N placebo). All graphs were produced with GraphPad Software.

For evaluation of sucrose preference, a mixed model factorial ANOVA was utilized where genotype, sex, and treatment were held as between-subject factors where the variable concentration of sucrose was held as a within-subjects factor. Regression analyses wee utilized to examine the concentration response curves. Similarly, a mixed model factorial ANOVA was used for evaluation of pre-pulse inhibition where transgene, sex, and treatment were held as between-subject factors where variable inter-stimulus interval was held as a within-subjects factor. To evaluate modified hole board, elevated-plus maze performance, and social behavior, a factorial ANOVA was employed to examine the effects of treatment, sex, and transgene. Rodent age was held as a covariate across all analyses.

Voltammetric recordings were obtained using customized software written in LabView (Knowmad Technologies LLC). Color plots of the evoked chemicals were generated within the data analysis features of the custom software. GraphPad Prism (version 5) was used to produce current versus time plots for each neurotransmitter of interest. Peak concentrations of neurotransmitter release were analyzed with a factorial ANOVA. Rates of release and reuptake of individual analytes were calculated using nonlinear regression where *K*, a nonlinear rate constant, was evaluated for both release and reuptake. Peak concentration values were obtained from the raw evoked electrical current.

Frequency distributions of spine parameters were compared using histograms of the entire data sets. Sholl analysis was performed using Neurolucida Explorer to examine dendritic branching and complexity. A mixed model ANOVA and discriminant function analysis were used to analyze spine parameters obtained from Sholl analysis.

## Results

### Sucrose Preference

Sucrose consumption curves plotted as a function of sucrose consumption are illustrated in **Figure 2** (**A-D**). Significant effects of genotype were observed on sucrose consumption (**Figure 2A**) and indicated by a genotype by concentration interaction [F(4, 224) =5.94, p≤0.0001] with a prominent quadratic component [F(1, 56) =6.18, p≤0.016]. Linear and non-linear modeling of the response curves to sucrose concentration further illustrated the HIV-1-mediated alteration. Specifically, dose-dependent responding to sucrose concentrations proceeded in a robust linear fashion (r^2^>0.97) in F344/N control animals whereas, in contrast, dose-dependent responding in HIV-1 Tg animals proceeded in a prominent quadratic fashion (one-phase association, r^2^>0.97). The difference in dose-response functions was not confounded with any positional or side bias of the animals as illustrated by mean sucrose consumption from each bottle position (**Figure 2B**). Differential effects of escitalopram treatment on the consumption curves were indicated between genotypes [F (4, 224) =5.94, p≤0.0001]. A prominent quadratic concentration by genotype by escitalopram interaction on was revealed [F(1, 56) =4.16, p≤0.046]. As shown, escitalopram did not alter sucrose consumption in the F344/N control animals; the dose-dependent sucrose consumption analysis revealed a global linear fit independent of escitalopram treatment (r^2^=0.92) (**Figure 2C**). However, for the HIV-1 Tg animals, escitalopram treatment restored the quadratic dose-response function (r^2^>0.97) to a prominent linear function (r^2^=0.88) (**Figure 2D**).

**Figure 2:**
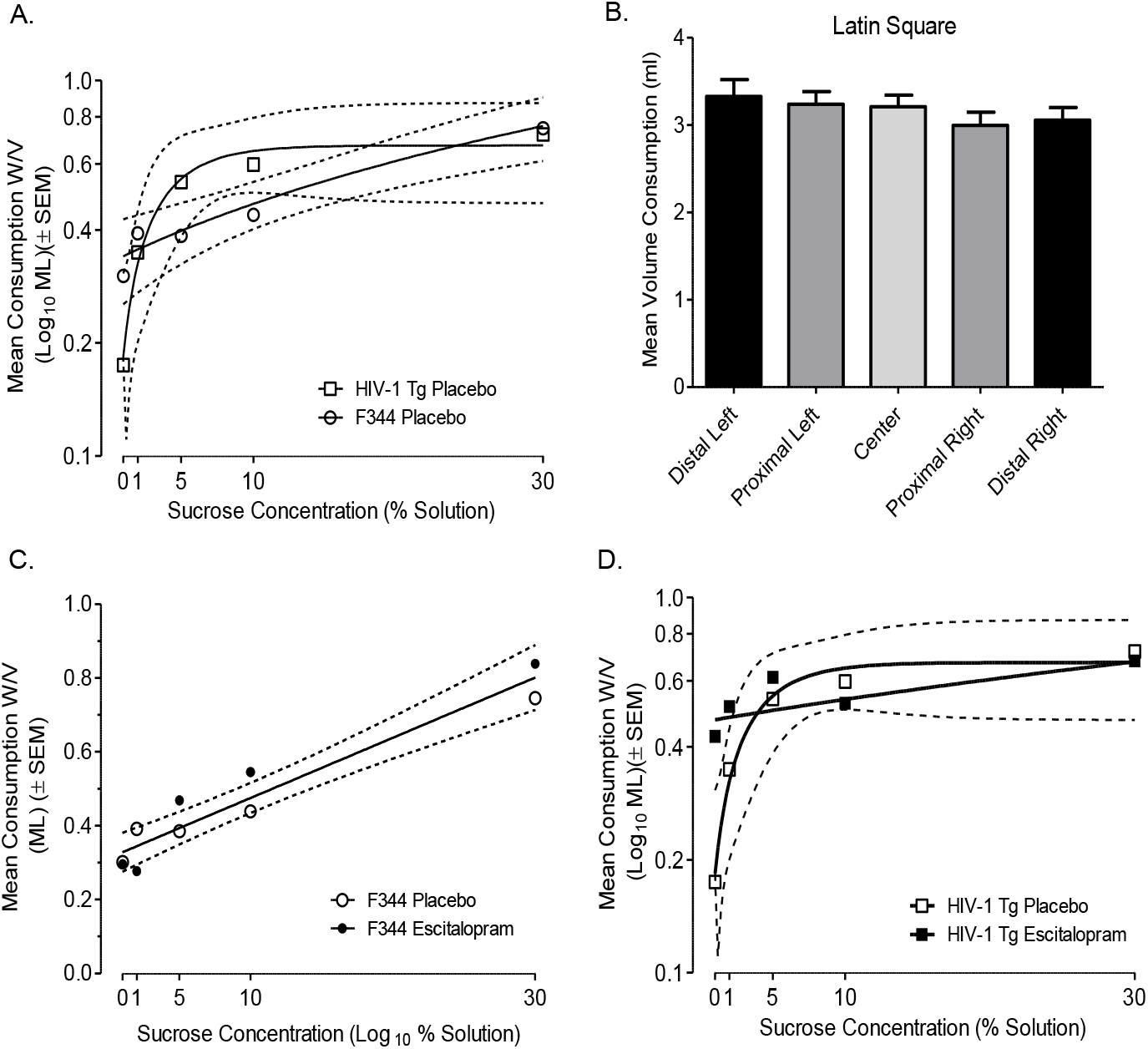
Sucrose preference testing. Five bottle sucrose preference test using a 0%, 1%, 5% 10% and 30% concentration. **A.** Dose-dependent responding to sucrose concentrations proceeded in a robust linear fashion (r^2^>0.97) in F344/N control animals whereas, in contrast, dose-dependent responding in HIV-1 Tg animals proceeded in a prominent quadratic fashion (one-phase association, r^2^>0.97). **B.** The difference in dose-response functions was not confounded with any positional or side bias of the animals as illustrated by mean sucrose consumption from each bottle position. **C.** Escitalopram did not alter sucrose consumption in the F344/N control animals; the dose-dependent sucrose consumption analysis revealed a global linear fit independent of escitalopram treatment (r^2^=0.92). **D.** However, for the HIV-1 Tg animals, escitalopram treatment restored the quadratic dose-response function (r^2^>0.97) to a prominent linear function (r^2^=0.88).

### Modified Hole Board

Exploration in the modified hole board (**Figure 3A**), as indexed by nose poke behavior, was sensitive to the effect of both genotype and escitalopram treatment [genotype by escitalopram interaction, F(1, 56) =12.18, p≤0.0009]. A significant effect of genotype was confirmed in the placebo treated animals [F(1, 56) =10.01, p≤0.0025] with the HIV-1 Tg animals displaying a greater number (>70%) of nose pokes than F344/N controls. It was also noted that there was no longer an effect of genotype after escitaolpram treatment [F(1,56)= 2.79, p≥0.10], Escitalopram treatment significantly decreased the number of nose pokes in HIV-1 Tg animals towards the levels of F344/N placebo controls [F(1, 56) =5.28, p≤0.025]. In contrast, escitalopram treatment increased the number of nose pokes in F344/N controls significantly above their placebo levels, an effect driven by the female animals [F(1, 56) =7.92, p≤0.007].

**Figure 3:**
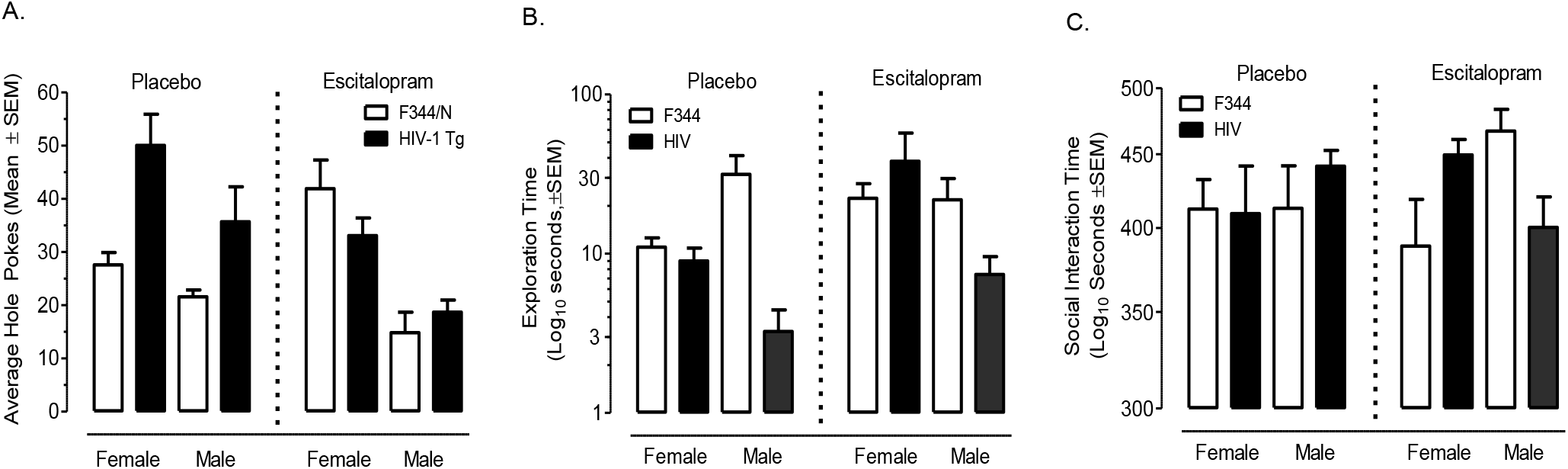
**A.** Exploration in the modified hole board, as indexed by nose poke behavior, was sensitive to the effect of both genotype and escitalopram treatment [genotype X escitalopram interaction, F(1, 56) =12.18, p≤0.0009]. **Left panel.** The HIV-1 Tg animals displaying a greater number (>70%) of nose pokes than F344/N controls [F(1, 56) =10.01, p≤0.0025]. **Right panel.** Escitalopram treatment significantly decreased the number of nose pokes in HIV-1 Tg animals towards the levels of F344/N placebo controls [F(1, 56) =5.28, p≤0.025]. In contrast, escitalopram treatment increased the number of nose pokes in F344/N controls significantly above their placebo levels, an effect driven by the female animals [F(1, 56) =7.92, p≤0.007]. **B.** Exploration time in the open arm of an elevated plus maze apparatus. **Left panel.** Exploration time in the elevated plus maze was not sensitive to an effect of the HIV-1 transgene [F(1, 72) =2.46, p>0.10], an effect clearly absent in females [F(1, 72) <1.0] but also not robust in the males [F(1, 72) =3.69, p=0.059]. **Right panel.** In the absence of a genotype effect, the evidence for an effect of escitalopram therapy was not expected nor observed. **C.** Social interaction time. Animals were tested with a novel sex and bodyweight matched partner in a novel cage across a ten minute trial period. **Left panel.** Total interaction time, recorded as the dependent measure of social behavior was insensitive to the effects of the HIV-1 transgene, escitalopram treatment, or their interaction [all Fs < 1.0]. **Right panel.** In the escitalopram treated animals [F(1,66)=6.67, p≤0.012] a genotype by sex effect was observed with M>F in F344/N vs F>M in HIV-1 Tg animals. The modulation of social behavior by escitalopram was not expected nor particularly relevant to its therapeutic potential given the insensitivity of the task to a genotype effect.

### Elevated Plus Maze

Exploration time in the elevated plus maze (**Figure 3B**) was less sensitive to an effect of the HIV-1 transgene [F(1, 72) =2.46, p>0.10], an effect clearly absent in both the females [F(1, 72) <1.0] but also not compelling in the male animals [F(1, 72) =3.69, p=0.059]. In the absence of a genotype effect, the evidence for an effect of escitalopram therapy was not expected nor observed. The overall increased exploration in female animals (independent of genotype) after repeated escitalopram treatment failed to meet statistical significance [F(1, 72) =3.01, p=0.087].

### Social and Play Behavior

Total interaction time, recorded as the dependent measure of social behavior (**Figure 3C**), was insensitive to the effects of the HIV-1 transgene, escitalopram treatment, or their interaction [all Fs < 1.0]. There was also no effect of sex, but a three-way interaction of genotype by sex by treatment was confirmed [F(1,66)=4.27, p≤0.043]. Interpretation of that latter term was guided by the significant two-way interaction of escitalopram treatment by sex in the F344/N animals [F(1,66)=4.26, p≤0.043] and of the genotype by sex two-way interaction in the escitalopram treated animals [F(1,66)=6.67, p≤0.012]. The presence of the differential genotype by sex effect is illustrated in the right panel of **Figure 3C** with M>F in F344/N vs F>M in HIV-1 Tg animals. The modulation of social behavior by escitalopram was not expected nor particularly relevant to its therapeutic potential given the insensitivity of the task to a genotype effect.

### Visual Pre-pulse Inhibition of the Acoustic Startle Response

Visual PPI was sensitive to both genotype and escitalopram treatment (**Figure 4A-4D**). As illustrated, a significant main effect of genotype [F(1,34)=10.26, p≤0.0029] accompanied by a prominent quadratic ISI by genotype interaction [F(1,34)=12.02, p≤0.0014] demonstrated a significant impairment in preattentive processing of the HIV-1 Tg animals relative to F344/N controls (**Figure 4A**). Further specific comparison of the ISI curves of the placebo treated animals of both genotypes revealed that the response amplitudes of the HIV-1 Tg animals were significantly less sensitive to response modulation by the ISI than F344/N controls [F(5,170)=5.31, p≤0.005]. There was no effect of sex nor interaction of sex with genotype or escitalopram or with ISI [Fs < 1.0] (**Figure 4B**). Escitalopram treatment did not appear to significantly alter the ISI curves of the F344/N animals (**Figure 4C**). In contrast, escitalopram treatment did significantly shift the shape of the ISI curves of the HIV-1 Tg animals toward that of the F344/N control animals, imparting a greater sensitivity to modulation by the ISI [F(5,170)=4.08, p≤0.02] (**Figure 4D**). Thus, in the presence of a significant genotype impairment in temporal sensitivity treatment with esticalopram was able to significantly restore functionality towards that of F344/N controls.

**Figure 4:**
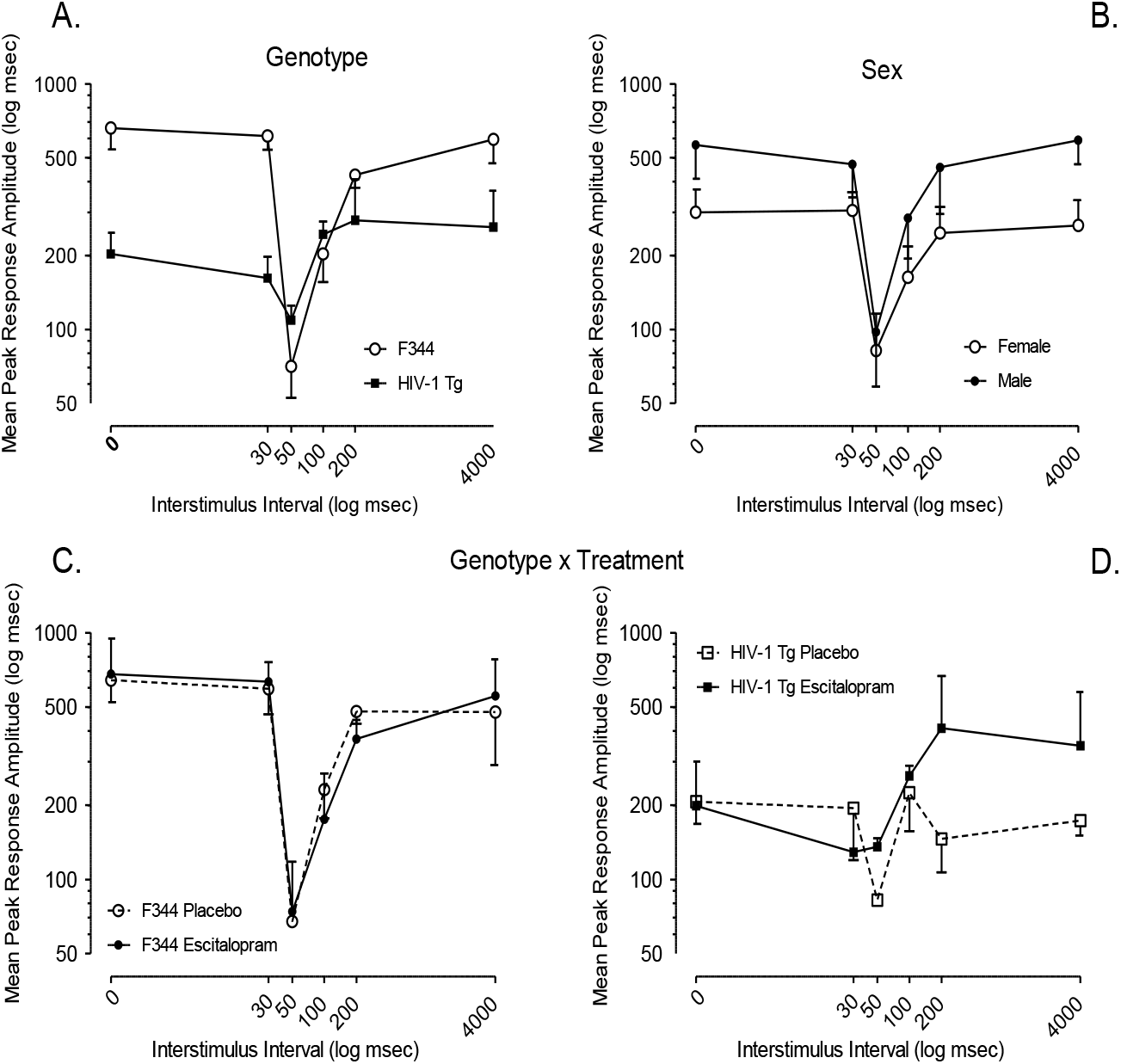
Visual Prepulse inhibition. Animals were tested during one session following a habituation session which occurred the day before. Visual PPI was sensitive to both genotype and escitalopram treatment. **A.** As illustrated, a significant main effect of genotype [F(1,34)=10.26, p≤0.0029] accompanied by a prominent quadratic ISI by genotype interaction [F(1,34)=12.02, p≤0.0014] demonstrated a significant impairment in preattentive processing of the HIV-1 Tg animals relative to F344/N controls, i.e., the response amplitudes of the HIV-1 Tg animals were significantly less sensitive to response modulation by the ISI than F344/N controls [F(5,170)=5.31, p≤0.005]. **B.** There was no effect of sex nor interaction of sex with genotype or escitalopram or with ISI [Fs < 1.0]. **C.** Escitalopram treatment did not appear to significantly alter the ISI curves of the F344/N animals. **D.** Escitalopram treatment did however significantly shift the shape of the ISI curves of the HIV-1 Tg animals toward that of the F344/N control animals, significantly restore functionality [F(5,170)=4.08, p≤0.02].

### Auditory Pre-pulse Inhibition of the Acoustic Startle Response

Auditory PPI was also sensitive to both genotype and escitalopram treatment (**Figure 5A-5D**). As illustrated, a significant main effect of genotype [F(1,37)=9.69, p≤0.0036] accompanied by a prominent quadratic ISI by genotype interaction [F(1,37)=8.97, p≤0.005] demonstrated a significant impairment in preattentive processing of the HIV-1 Tg animals relative to F344/N controls (**Figure 5A**). Further specific comparison of the ISI curves of the placebo treated animals of both genotypes revealed that the response amplitudes of the HIV-1 Tg animals were significantly less sensitive to response modulation by the ISI than F344/N controls [F(5,170) =5.31, p≤0.008]. There was no effect of sex nor interaction of sex with genotype or escitalopram or with ISI [Fs < 2.0] (**Figure 5B**). Escitalopram treatment did not appear to significantly alter the ISI curves of the F344/N animals (**Figure 5C**). Although escitalopram treatment appeared to shift the shape of the ISI curves of the HIV-1 Tg animals toward that of the F344/N control animals (**Figure 5D**; peak inflection away from 200 msec), a significant difference between the ISI curves remained between the two genotype groups [F(5,185) =3.87, p≤0.038]. Overall, in the presence of a significant genotype impairment in temporal sensitivity esticalopram was able to facilitate a partial restoration of functionality towards that of F344/N controls.

**Figure 5:**
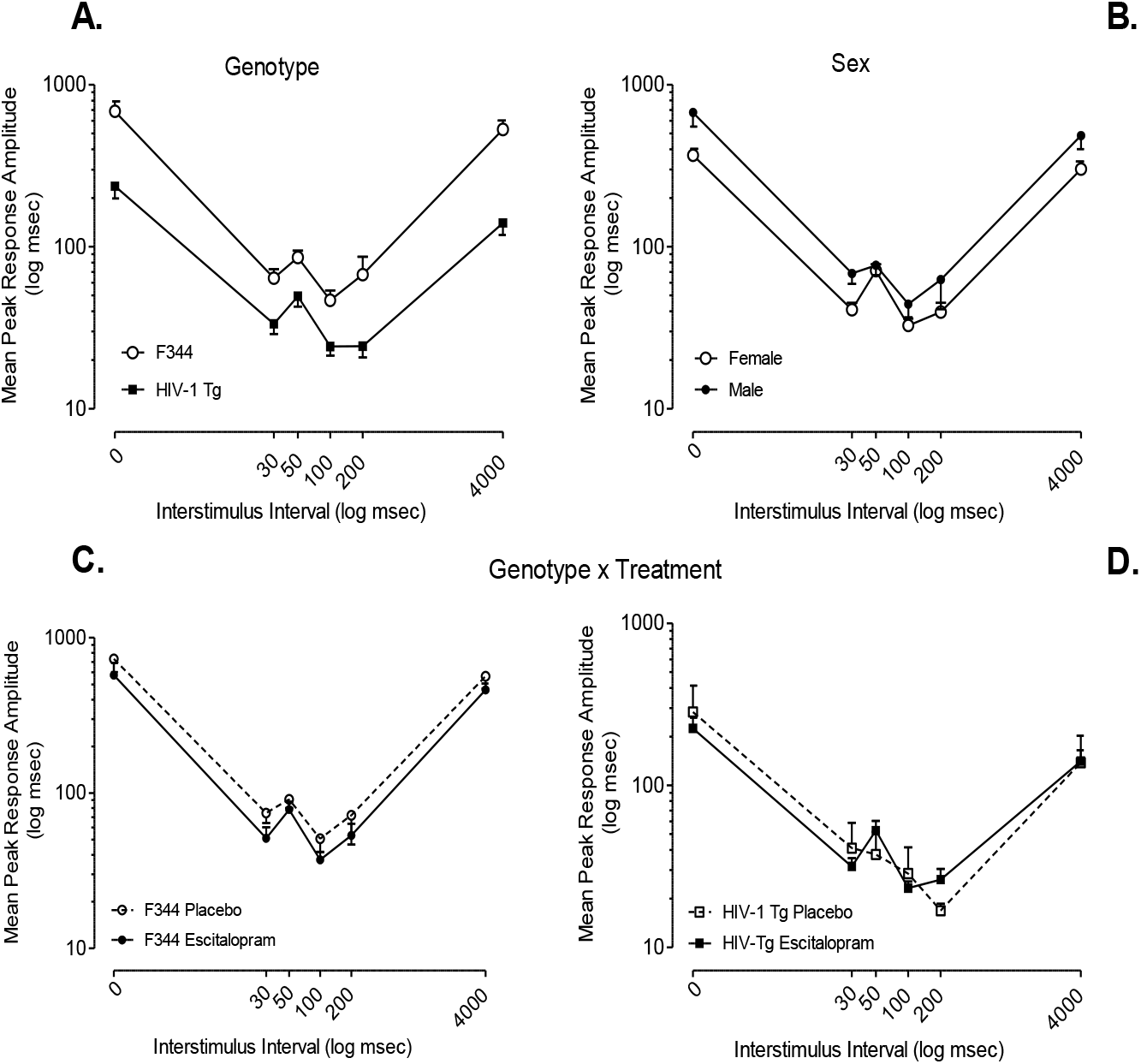
Auditory Prepulse inhibition Animals were tested during one session following a habituation session which occurred the day before. Auditory PPI was also sensitive to both genotype and escitalopram treatment. **A.** As illustrated, a significant main effect of genotype [F(1,37)=9.69, p≤0.0036] accompanied by a prominent quadratic ISI by genotype interaction [F(1,37)=8.97, p≤0.005] demonstrated a significant impairment in preattentive processing of the HIV-1 Tg animals relative to F344/N controls, i.e., the response amplitudes of the HIV-1 Tg animals were significantly less sensitive to response modulation by the ISI than F344/N controls [F(5,170) =5.31, p≤0.008]. **B.** There was no effect of sex nor interaction of sex with genotype or escitalopram or with ISI [Fs < 2.0]. **C.** Escitalopram treatment did not appear to significantly alter the ISI curves of the F344/N animals. **D.** Although escitalopram treatment appeared to shift the shape of the ISI curves of the HIV-1 Tg animals toward that of the F344/N control animals (peak inflection away from 200 msec) restoring functionality, a significant difference did remain between the two genotype ISI curves [F(5,185) =3.87, p≤0.038].

### Dopamine and Serotonin Voltammetry

Decreases in peak transmission and reuptake of dopamine (**Figure 6A-C**) and serotonin (**Figure 7A-C**) were found in the HIV-1 Tg rat. Maximal evoked concentration (Cmax) was impaired in transgenic animals across both dopamine and serotonin recordings. [Dopamine, F(1,9)=33.25, p≤0.001; Serotonin, F(1,16)=60.97, p≤0.001]. Additionally, rates of reuptake as defined by the nonlinear rate constant (*k*) were impaired in transgenic animals relative to control animals (Denton, 2019). [Dopamine, F344/N *K=0.43*, HIV-1Tg *K=0.73* F(1,2634)=19.19, p≤0.001; Serotonin, F344/N *K=0.37*, HIV-1Tg *K=0.56* F(1,4314)=7.308, p≤0.05.]

**Figure 6.**
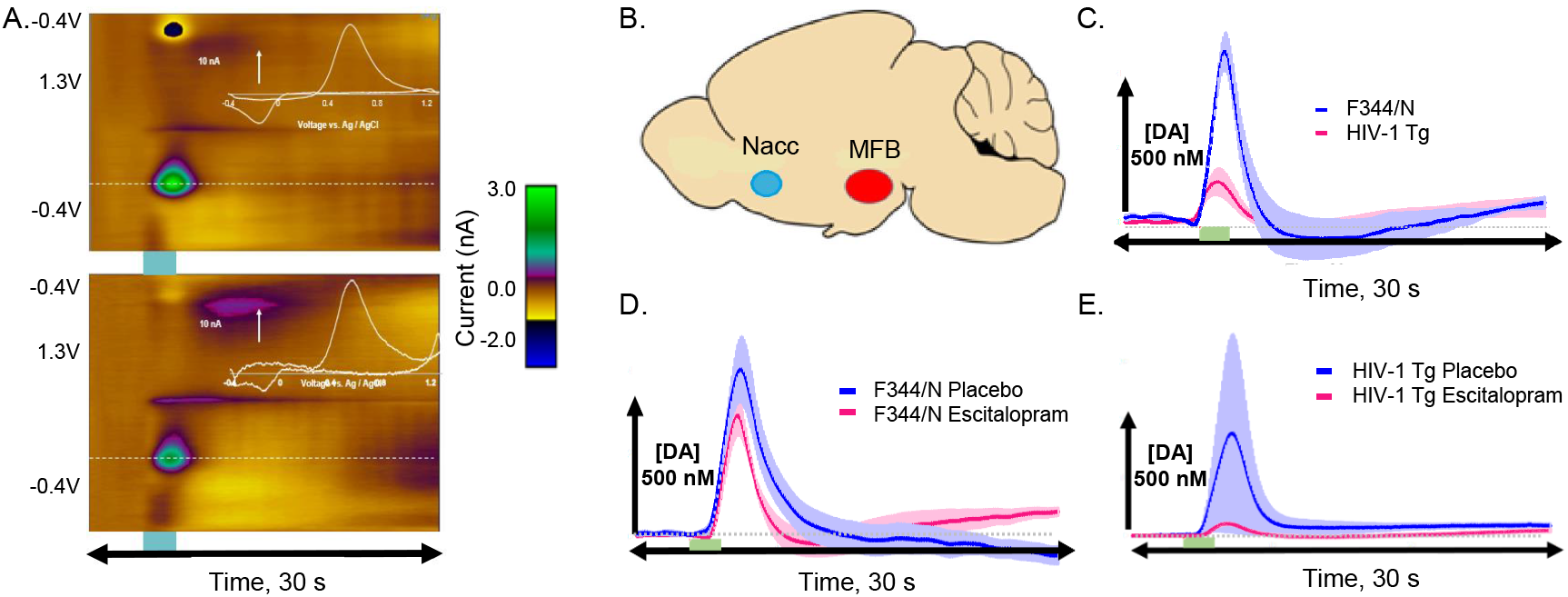
Fast Scan Cyclic Voltammetry of dopamine **A**: Colorplots for HIV-1 Tg animals (top) and F344/N animals (bottom). **B.** Stimulus pin implant location (red) and recording carbon fiber microelectrode (blue). **C.** Evoked dopaminergic potentials for HIV-1 Tg animals (pink) and F344/N controls (blue). Dopamine peak release was impaired in HIV-1 Tg animals relative to F344 controls [F(1,9)=33.25, p≤0.001]. Additionally, rates of reuptake were significantly impaired in the HIV-1 Tg rat [F344/N *K=0.43*, HIV-1Tg *K=0.73* F(1,2634)=19.19, p≤0.01]. **D.** Evoked dopaminergic potentials for F344/N animals treated with placebo (blue) and F344/N animals treated with escitalopram (pink). **E.** Evoked dopaminergic potentials for HIV-1 Tg animals treated with placebo (blue) and HIV-1 Tg animals treated with escitalopram (pink). Escitalopram treatment did not attenuate dopaminergic release deficits in HIV Tg animals [F(1,18)=0.123, p=ns.]. Moreover, rates of reuptake were not significantly different for HIV-1 animals treated with escitalopram [(*k=0.41* F(1,1414)=0.47, p=ns].

**Figure 7.**
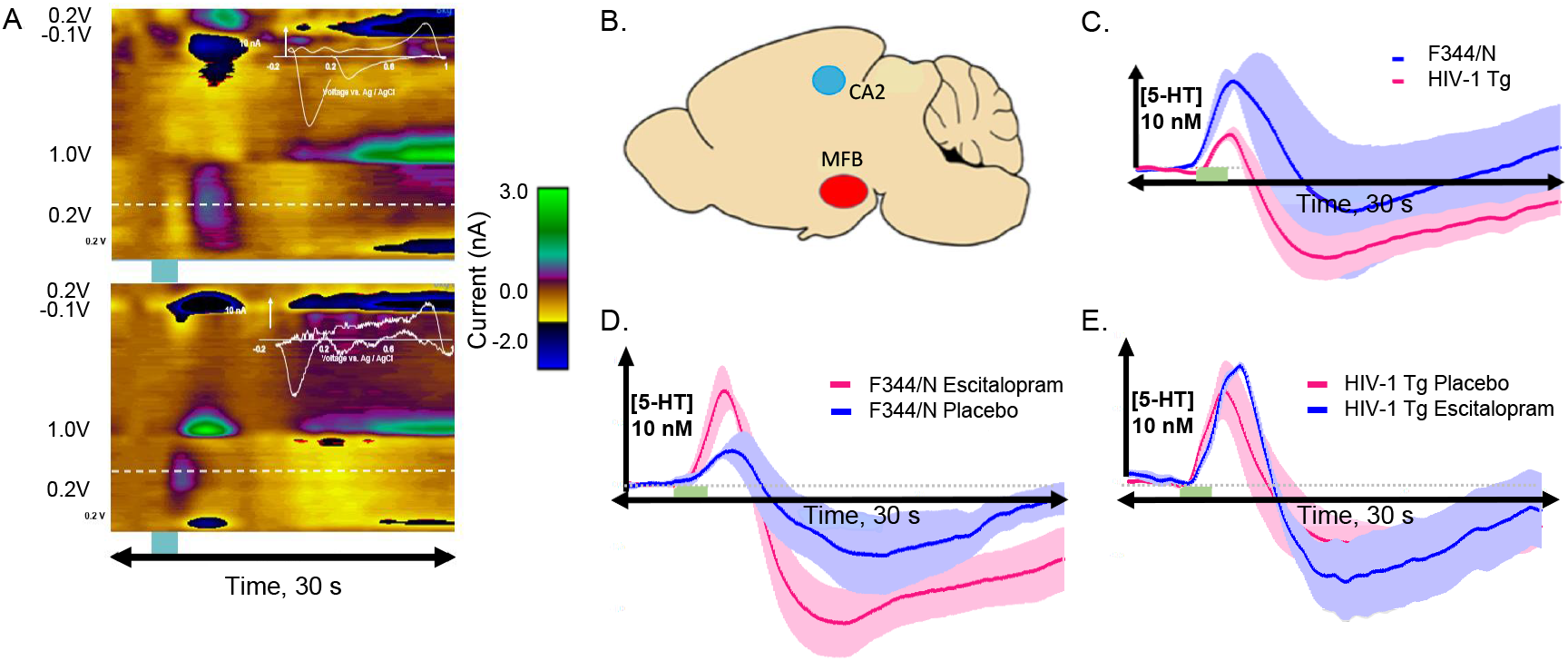
Fast Scan Cyclic Voltammetry of serotonin **A**: Colorplots for HIV-1 Tg animals (top) and F344/N animals (bottom). **B.** Stimulus pin implant location (red) and recording carbon fiber microelectrode (blue). **C.** Evoked serotonergic potentials for HIV-1 Tg animals (pink) and F344/N controls (blue). Serotonin peak release was impaired in HIV-1 Tg animals relative to F344 controls F(1,16)=60.97, p≤0.001]. Additionally, rates of reuptake were significantly impaired in the HIV-1 Tg rat [F344/N *K=0.37*, HIV-1Tg *K=0.56* F(1,4314)=7.308, p≤0.05.] **D.** Evoked serotonergic potentials for F344/N animals treated with placebo (blue) and F344/N animals treated with escitalopram (pink). **E.** Evoked serotonergic potentials for HIV-1 Tg animals treated with placebo (blue) and HIV-1 Tg animals treated with escitalopram (pink). Escitalopram treatment did not attenuate serotonin release deficits in HIV Tg animals, but did marginally improve peak serotonin concentration in F344 animals [F(1,30)=0.99, p=ns]. Rates of reuptake were slower for animals treated with escitalopram than for placebo-treated animals [*k=0.32* vs. *k=0.20* F(1,7674)=23.75, p≤0.01].

No statistically significant differences relative to control were found for dopamine transmission in HIV-1 Tg rats treated with escitalopram [F(91,150)=1.00, p=≥ 0.05 (**Figure 6D-E**). Evoked rates of maximal dopamine release were not statistically significant across a genotype by treatment analysis [F(1,18)=0.123, p=≥ 0.05.], though HIV-animals treated with SSRI medication demonstrated the lowest peak concentration. Rates of reuptake were not statistically different for HIV-1 Tg animals treated with escitalopram, compared to control [*(k=0.41* F(1,1414)=0.47, p ≥ 0.05], though F344/N animals treated with escitalopram demonstrated slower rates of reuptake than animals treated with placebo [*k=0.49* vs. *k=0.23* F(1,3834)=16.1, p≤0.001] (Denton, 2019).

Increases in serotonin transmission were found in F344/N control animals treated with escitalopram, but not in HIV-1 Tg rodents (**Figure 7D-E**). Rates of clearance (reuptake) were slower in animals treated with escitalopram (*k=0.55*) relative to animals treated with placebo (*k=0.34*) [F(1,4074)=9.18, p≤0.05]. While F344/N animals treated with escitalopram displayed a 55% increase in peak evoked serotonergic potential, the effect was not significant [F(1,30)=0.99, p=ns] although rates of reuptake were altered for animals treated with escitalopram [*k=0.32* vs. *k=0.20* F(1,7674)=23.75, p≤0.01].

### Medium Spiny Neuron Branching/Morphology

Overall, MSNs of HIV-1 Tg animals treated with escitalopram exhibited greater dendritic length, volume, and intersections at distal radii, demonstrating that escitalopram was effective in promoting dendritic complexity and proliferation in the nucleus accumbens of HIV-1 Tg animals. Frequency distributions of spine length of medium spiny neurons in the nucleus accumbens revealed a genotype/treatment interaction effect with escitalopram altering length distributions for both HIV-1 Tg and F344/N animals (**Figure 8A-B**). However, escitalopram did not appear to alter frequency distributions for head diameter or volume. Moreover, escitalopram appeared to alter spine morphology in HIV-1 Tg rats, as individuals treated with escitalopram exhibited higher frequencies of stubby and mushroom spine types across successive radii when compared with placebo-treated HIV-1 Tg animals (**Figure 8C-D**). Sholl analysis revealed a statistically significant interaction effect for treatment and genotype upon dendritic proliferation. HIV-1 Tg animals demonstrated markedly less dendritic complexity when compared with control counterparts. However, treatment with escitalopram served to dramatically improve dendritic complexity in HIV-1 Tg animals, even normalizing animals to control levels with respect to dendritic intersections at concentric radii (**Figure 9A-B**). [F(1,8)=5.34, p<0.05]. A similar effect was found with respect to length and volume, although the effects were not statistically significant [F(1,13)=2.18, p≥ 0.05; F(1,13)=1.51, p≥0.05, respectively]. Additionally, a discriminant function analysis with jackknife resampling procedure was performed to determine whether animals could be accurately classified into treatment groups (placebo vs. escitalopram) based upon dendrite intersections obtained from the Sholl analysis. Using the parameter of dendrite intersection/radii for each of the concentric radii, animals were correctly classified into treatment groups with 100% accuracy [Wilks’ λ=0.216, x ^2^_(12)_=26.02, p≤0.05].

**Figure 8.**
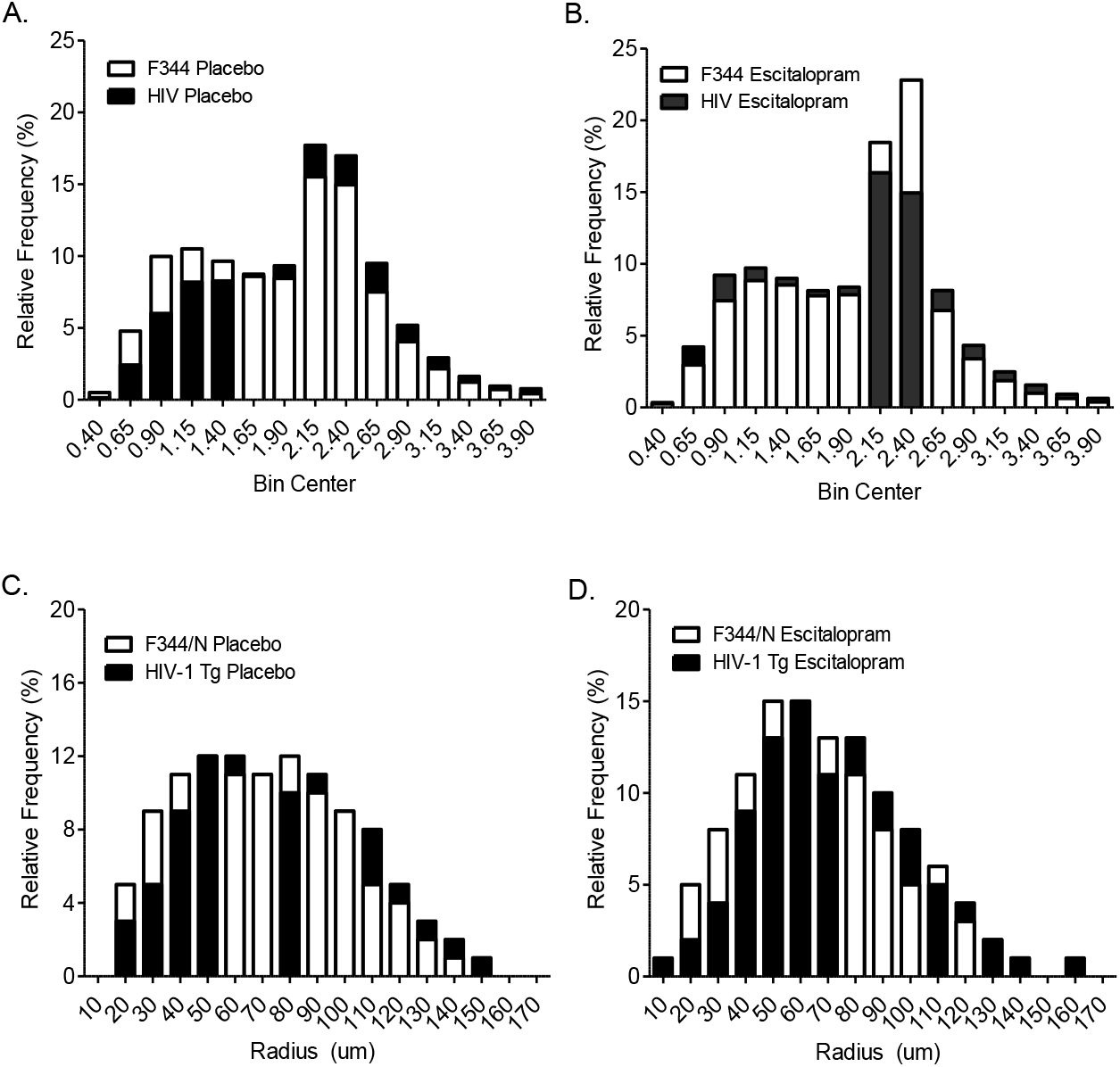
Chronic escitalopram treatment produced increased populations of mushroom spines. **A:** Relative frequency distributions of dendrite length of MSNs in the nucleus accumbens are illustrated for placebo controls as a function of genotype; note the leftward shift in the HIV-1 Tg animals. **B.** Relative frequency distributions of dendrite length of MSNs in the nucleus accumbens are illustrated for esticalopram treated animals as a function of genotype; note the reduction in peakedness in the HIV-1 Tg animals. **C-D.** Frequency distributions of mushroom dendritic spines displayed across concentric radii also showing shifts as a function of genotype (**C**) and esticalopram (**D**).

**Figure 9.**
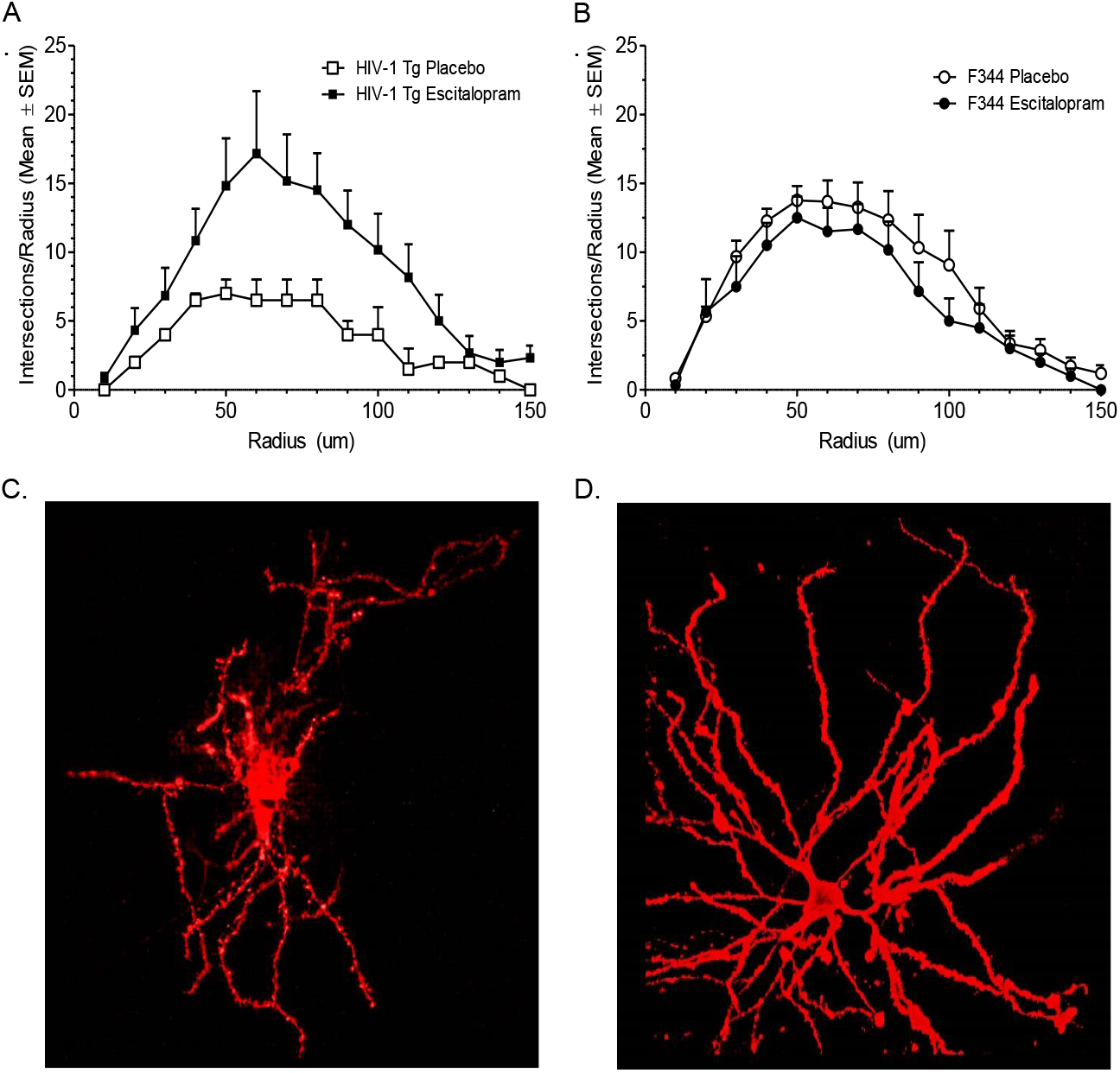
Escitalopram treatment dramatically increased dendritic proliferation and connectivity in HIV-1 Tg rats. **A:** Sholl analysis of dendritic intersections/radii are shown for HIV-1 Tg animals separated by treatment condition. **B.** Sholl analysis of dendritic intersections/radii are shown for F344/N animals separated by treatment condition. Treatment with escitalopram served to dramatically improve dendritic intersections at distal radii in HIV-1 Tg animals [F(1,8)=5.34, p<0.05], normalizing dendritic complexity similar to control levels. Confocal images of medium spiny neurons from the nucleus accumbens are presented for HIV-1 Tg rats treated with placebo (**C**) and escitalopram (**D**).

## Discussion

Chronic escitalopram treatment significantly increased dendritic complexity and altered spine morphology in the HIV-1 Tg rat. Previous reports found extensive HIV-induced damage to MSNs in the nucleus accumbens of HIV-1 Tg rats (Roscoe et al., 2014; McLaurin et al., 2018) thus, synaptodendritic restoration may be a key target for therapeutic intervention. We found that chronic escitalopram administration was successful in restoring dendritic complexity to MSNs in the nucleus accumbens of HIV-1 Tg rats, even to control levels. These findings suggest therapeutic efficacy for escitalopram in repairing HIV-mediated damage in the nucleus accumbens. The behavioral and preattentive processing tasks that were sensitive to impairments in the HIV-1 Tg animals were also sensitive to demonstrating functional improvements with escitalopram treatment. However, escitalopram treatment did not restore neurotransmission deficits in HIV-1 Tg rats. Serotoninergic functioning in F344 animals was improved by escitalopram treatment; in contrast, HIV-1 Tg animals treated with escitalopram failed to display an increase in serotonergic functioning. Nevertheless, escitalopram treatment represents an important first step toward effective therapeutic intervention in repairing HIV-mediated synaptodendritic damage and restoring functional impairments following exposure to HIV-1 proteins. A summary of the observed effects is included in Figure 10.

**Figure 10.**
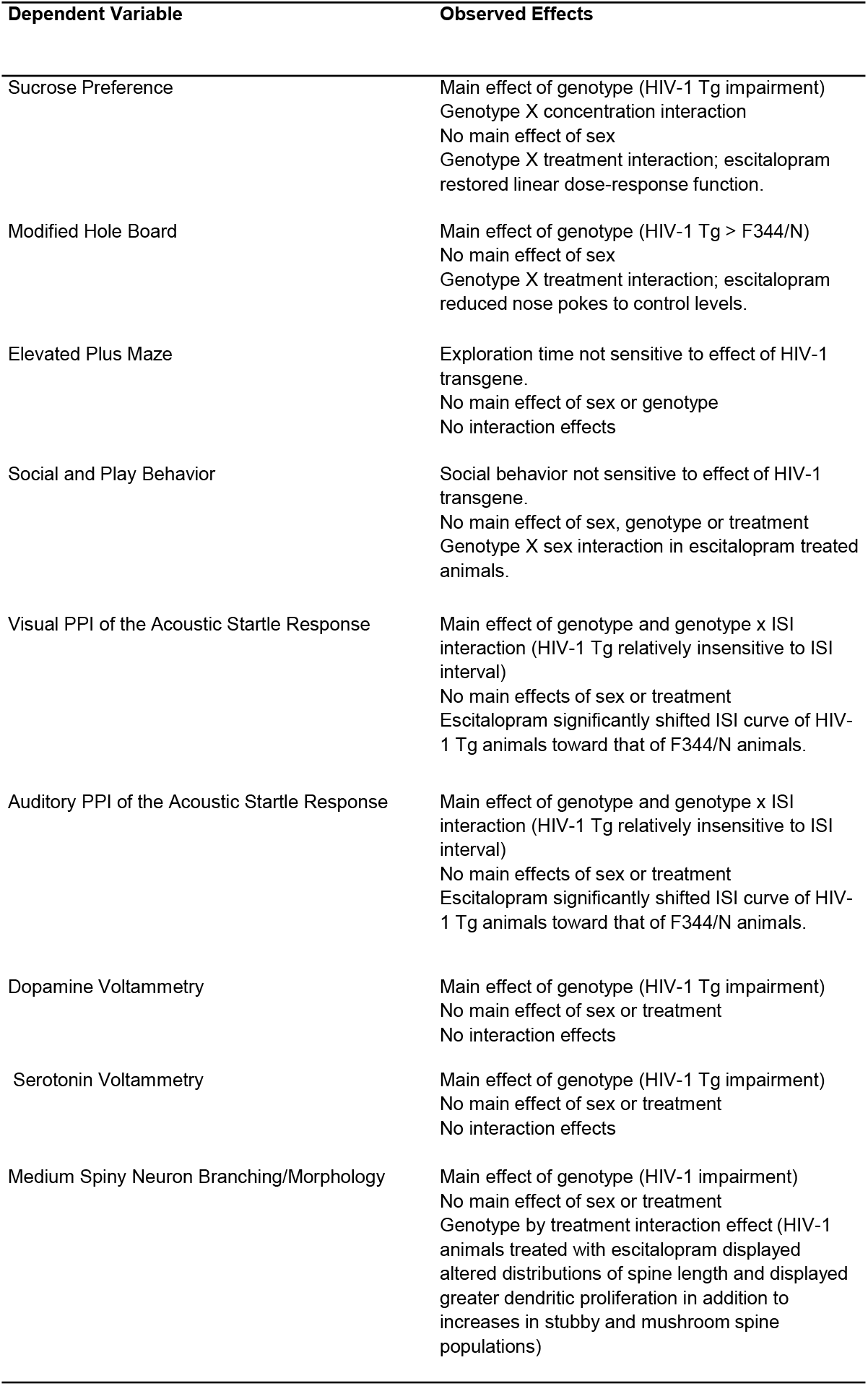
Summary of observed effects.

Simultaneous decreases in release and reuptake rates for dopamine and serotonin activity were found in HIV-1 Tg rats, relative to F344/N controls. Dopaminergic impairments in the nucleus accumbens are consistent with previous research (Javadi-Paydar et al. 2017), which employed *ex vivo* striatal brain slices to examine DA reuptake in HIV-1 Tg rats. Moreover, the present findings are consistent with previous reports (Denton et al., 2019), which demonstrated severe monoamine dysfunction in intact HIV-1 Tg rats. Specifically, dopaminergic functioning in the HIV-1 Tg rat following long-term HIV-1 protein exposure is characterized by slower reuptake rates from diminished peak concentrations of dopamine. (Denton et al., 2019). These findings are consistent with both preclinical PET imaging studies (Sinharay et al. 2017) and clinical PET imaging studies (Chang et al. 2008). Acute in vivo studies (Ferris et al., 2009) reported decreased extracellular striatal dopamine concentrations following HIV-1 Tat protein infusion. In contrast, others have reported transitory increases in dopamine levels in the nucleus accumbens (at 3 days; Kesby et al., 2016a) which did not persist (absent at 10 days; Kesby et al., 2016b), following acute induction of Tat protein expression in astrocytes (Kesby et al., 2017). The inconsistency of dopamine levels following acute Tat protein exposure is further illustrated by the puzzling findings of increased dopamine concentrations in the prefrontal cortex at 7 days post-induction of Tat, without any accompanying changes in striatal dopamine levels (Strauss et al., 2020). Although it is difficult to reconcile findings from acute studies (i.e, 1-2 weeks) in terms of dopaminergic alterations following HIV-1 protein exposure (i.e., reports of regionally specific increases, decreases, and no changes in dopamine), with findings from chronic HIV-1 protein exposure (months to years via the HIV-1 Tg rat, i.e., similar to long-term HIV-1 exposure in humans) which clearly and reliably show decreased extracellular dopamine levels and impairs mesocorticolimbic circuit neurotransmission (for review, see Illenberger et al., 2020; Bertrand et al., 2018). The decreases in dopaminergic functioning play a key role in HIV-1 apathy (Bertrand et al., 2018) and HAND (Moran et al., 2019; McLaurin et al., 2020).

The nucleus accumbens core and shell regions receive dense input from serotonergic neurons from the raphe nucleus (Van Bockstaele et al., 1993). The shell region of the nucleus accumbens contains axons and axon terminals that are of larger diameter and greater varicosity than those observed in the nucleus accumbens core region. Axon terminals in the nucleus accumbens core contain vesicles of greater density and form symmetrical synaptic contact with dendrites in this region. Consequently, serotonin has been reported to regulate dopamine release within the nucleus accumbens in both an excitatory and inhibitory manner (Parsons et al., 1993). Serotonin has been shown to increase dialysate dopamine in the nucleus accumbens (Parsons et al., 1993), whereas 5-HT_2C_ receptor agonists have been shown to decrease mesocorticolimbic dopamine function, with 5-HT_2C_ antagonists showing the opposite effect (Di Matteo et al., 2001). Thus, interactions between serotonin and dopamine might be anticipated in the HIV-1 Tg rat, and in these studies both neurochemical systems reacted similarly to long-term HIV-1 protein exposure.

Presently, we report no sex differences in dopaminergic transmission between male and female rats. These findings are in line with previous studies from our laboratory employing fast-scan cyclic voltammetry which did not elucidate an effect of sex in dopaminergic release or reuptake (Denton et al., 2019). Moreover, these findings are in line with findings from microdialysis studies examining sex differences in dopamine between male and female rats (Egenrieder et al., 2020). In a review and meta-analysis, Egenrieder and colleagues (2020) report that there are consistently no sex differences in dopamine levels in either the caudate putamen or nucleus accumbens. In the current studies, female animals were sacrificed on the day of diestrus; an estrus stage of low ovarian hormone levels. Our prior studies (Bertrand et al., 2018) were conducted in ovariectomized female animals. Future studies may study the role of ovarian hormones across the estrus cycle to determine how estrogen and progesterone levels may impact dopaminergic and serotonergic function in the HIV-1 Tg rat.

Serotonin transmission using FSCV is characterized by a quick rise to peak evoked concentration followed by reuptake mechanisms specific to the site being measured (West et al., 2018; Abdalla et al., 2019). These site-specific uptake phases were identified as Uptake 1 and Uptake 2 (Shaskan and Snyder 1970). Uptake 1 occurs as a result of the activity of the (SERT). Consequently, Uptake 1 is characterized by a high affinity for serotonin molecules but low efficiency. Studies of serotonin recordings in mice have characterized Uptake 1 as a single decay curve from peak concentration that is relatively slow and persists for approximately 12 seconds. Uptake 2 occurs as a result of non-SERT transporters and is characterized by higher efficiency, but lower capacity and affinity. Studies in mice have shown that Uptake 2 has a much shorter decay curve that reaches baseline activity quickly (Abdalla et al., 2019). These studies have further suggested that serotonin release in the hippocampus is largely mediated by Uptake 2 (non-SERT) mechanisms (West et al., 2018; Abdalla et al., 2019). Such findings may potentially explain why escitalopram was not found to alter serotonin kinetics in HIV-1 Tg animals in the present study. Escitalopram is known for its high affinity for SERT (Braestrup and Sanchez, 2004), but as the kinetics of hippocampal serotonin do not appear to be as highly mediated by SERT, the present findings may result from escitalopram failing to mediate hippocampal specific serotonin kinetics, such as Uptake 1. Future experiments examining prefrontal cortex kinetics using FSCV, (previous research from our lab has indicated synaptodendritic and functional impairment in the PFC – McLaurin et al., 2018; Denton et al., 2019), would be informative relative to the role of SERT in HIV-1.

Synaptic loss, without neuronal death, is associated with HIV-1 and likely underlies neurocognitive impairments (Everall et al., 1999; Ellis et al., 2009). As HIV-1 does not directly infect neurons, synaptic loss is a result of exposure to viral products such as HIV Tat and gp120 (Fitting et al., 2015). Specifically, the cysteine-rich region of the Tat protein has been shown to play a critical role in the development of synaptic loss (Bertrand et al., 2013). Synaptic damage may be a result of Tat-produced proteasome-mediated degradation of micro-tubule-associated protein 2 (MAP2), which consequentially results in a collapse of cytoskeletal filaments and spine/synaptic loss (Kim et al., 2008). Whereas cellular death requires calcium-mediated neuronal nitric oxide synthesis, synaptic damage associated with Tat is not mediated by nNOS, but rather ubiquitin-proteasomal pathways (Aprea et al., 2006; Kim et al., 2008). HIV-induced synaptic loss may result from a compensatory process to avoid cellular death (Green et al., 2019), thereby altering circuit connectivity (Illenberger et al., 2020). HIV viral proteins and inflammatory cytokines in the brain result in excessive activation of glutamatergic pathways, particularly in the frontostriatal pathways, which are critical for apathy. Previous reports highlight the potential reversibility of HIV-1 induced dendritic damage, including the present report (Kim et al., 2008; Kim et al., 2011; Bertrand et al., 2014). More research is needed to more fully elucidate effective treatments for HIV-1 induced synaptodendritic damage, although phytoestrogen treatment (Bertrand et al., 2014; McLaurin et al., 2020), cannabinoid receptor activation (Kim et al., 2011), and the presently discussed SSRI treatment are promising treatment avenues. Moreover, functional endpoints of neurocognition are essential (McLaurin et al., 2019) in any assessments of neurorestorative treatments for apathy and HAND.

What is unclear from the present findings, however, is why escitalopram-mediated improvement in the dendritic spine profile did not produce a likewise improvement in neurochemistry, particularly regarding dopaminergic functioning in the nucleus accumbens. The present finding that escitalopram treatment increased mushroom spine proliferation, suggests the potential to rectify many of the deleterious effects of HIV-1. Mushroom spines are associated with the upper limits of synaptic strength and represent mature synaptic connections (Yuste, 2010; Berry & Nedivi, 2017). Moreover, these spine subtypes have the largest spine head volume of all spine subtypes, which correlates to a larger pre-synaptic zone and post-synaptic density. Increased size of the post-synaptic density is furthermore correlated with both the size of the active pre-synaptic zone and the number of docked pre-synaptic vesicles (Berry & Nedivi, 2017). Escitalopram treatment also increased in stubby spine subtypes. The stubby spine types may have little to offer in the context of neurotransmission, as their shape and lack of a voluminous head do not engender effective neuronal communication when compared to spine subtypes such as thin or mushroom (Yuste, 2010; Bae et al., 2012). Though stubby spine populations are maintained in the adult brain, they are typically considered to be markers of incomplete synaptic development (Yuste, 2010; Berry & Nedivi, 2017). Moreover, increases in stubby spine proliferation are often associated with neuropathology, including clinical depression (Buyukdura et al., 2013). The findings that escitalopram increases both mushroom and stubby subtypes suggest that escitalopram treatment may be a first step toward repair of HIV-mediated damage in the nucleus accumbens.

The most likely explanation for why an increase in neurotransmission was not observed is the timing of the present investigations, coupled with the likelihood that full restoration of neurochemical processes and circuitry occurs slower than the formation of dendritic spines. Such a hypothesis would explain why escitalopram produced an increase in spine subtypes conventionally associated with immaturity while likewise increasing populations of those types that represent the upper limits of synaptic strength. Future studies of chronic escitalopram may show increased neurotransmission if measured at longer times from treatment onset. Thus, while escitalopram has the potential to dramatically increase dendritic branching and spine proliferation within six weeks of treatment, SSRI therapies may require a longer treatment period to reach full effect. Nevertheless, spine loss in the context of HIV-1 can be recovered by therapeutic intervention. Full recovery of circuit potential may take longer than changes in dendritic spines, thus a longer investigation may observe the full restoration of circuit connectivity and attenuation of dopaminergic and serotonergic deficits following treatment with escitalopram.

Visual and auditory PPI impairments in the HIV-1 Tg rat have been consistently reported by our laboratory (Moran et al., 2013; McLaurin et al., 2016; McLaurin et al., 2017; McLaurin et al., 2018). SSRI treatment was effective in restoring function for both visual or auditory PPI deficits in the HIV-1 Tg rat, albeit the evidence was stronger for the visual PPI task. HIV-1 Tg animals displayed consistent abnormalities in PPI across interstimulus interval (ISI). The additional behavioral tasks that were sensitive to the HIV-1 transgene, e.g., sucrose preference and hole board, both also displayed convincing evidence of restoration of function with escitalopram treatment. The curvilinear shift in response to variable sucrose concentration observed for HIV-Tg rats was restored to a linear function with escitalopram. The observed curvilinear shift in sucrose concentration responses for HIV-1 Tg rats is a novel finding within the literature, as previous reports have not indicated a significant difference between sucrose response in HIV-Tg and F344 animals (Bertrand et al., 2018); however, that prior report was conducted with ovariectomized female animals. Rodent performance in the modified hole board revealed a statistically significant effect of genotype, with a prominent increase in nose poke behavior of the HIV-1 Tg animals. Escitalopram treatment significantly reduced that exploratory behavior to approximate F344/N control levels. Neither the plus maze nor social behavior tasks were particularly sensitive to any impairment by the HIV-1 transgene; given this insensitivity no specific therapeutic effect of escitatolpram would be expected. Overall, escitalopram was quite effective at modulating behavioral responses in HIV-1 Tg rats in each and every task that was sensitive to detecting HIV-1 protein induced impairments, suggesting escitalopram enabled functional recovery from HIV-1.

## Conclusions

In the present study, therapeutic efficacy of escitalopram was found in treating HIV-1 Tg rats when examining spine complexity, as SSRI treatment was found to increase dendritic proliferation in HIV-1 Tg rats and consequently normalize these animals to control levels of complexity and proliferation. Similarly, the therapeutic efficacy of esticalopram was suggested in each of the behavioral tasks that were sensitive to impairments produced by the HIV-1 transgene. Given that the present findings reveal that escitalopram did not improve serotonergic tone in the HIV-1 Tg rat, but increased transmission in control rats, it is likely the case that dendritic repair precedes restoration of circuit neurotransmission and function. In sum, chronic escitalopram treatment is an effective therapeutic approach for HIV-1 mediated synaptodendritic damage as well as for HIV-1 induced behavioral impairments, and moreover, has the potential for repair of HIV-1 neurological deficits and functional restoration of HAND.

## Compliance with Ethical Standards

This experiment was conduction in accordance with the recommendations of the National Institute of Health’s Guide for the Care and Use of Laboratory Animals. Research protocols used were approved by the University of South Carolina Institutional Animal Care and Use Committee (assurance number: D16-00028). Additionally, the authors report no conflicts of interest or competing financial interests.

## Acknowledgments

This research was supported by National Institute of Health Grants NS100624, DA013137, HD043680, MH106392 & by a National Institute of Health T32 Training Grant 5T32GM081740. The authors acknowledge and thank Dr. Michael Cranston and Dr. Hailong Li for their assistance with this project.

## References

Abdalla A, West A, Jin Y, Saylor R, Qiang B, Pena E, (…) Hashemi P (2019) Fast serotonin voltammetry as a versatile tool for mapping dynamic tissue architecture: I. Responses at carbon fibers describe local tissue physiology. J. Neurochem doi:10.1111/jnc.14854

Aksenov MY, Aksenova MV, Silvers JM, Mactutus CF, Booze RM (2008) Different effects of selective dopamine uptake inhibitors GBR 12909 and WIN 35428 on HIV-1 Tat toxicity in rat fetal midbrain neurons. Neurotoxicology 29:971–977

Aprea S, Del Valle L, Mameli G, Sawaya BE, Khalili K, Peruzzi F (2006) Tubulin-mediated binding of human immunodeficiency virus-1 Tat to the cytoskeleton causes proteasomal-dependent degradation of microtubule-associated protein 2 and neuronal damage. J Neurosci 26:4054–4062.

Arseniou S, Arvaniti A, Samakouri M (2014) HIV infection and depression. Psychiatry Clin Neurosci 68:96–109

Bae J, Sung BH, Cho IH, Kim SM, Song WK. (2012) NESH regulates dendritic spine morphology and synapse formation. PLoS One 7:e34677

Berry KP, Nedivi E (2017). Spine dynamics: Are they all the same? Neuron, 96, doi:10.1016/j.neuron.2017.08.008.

Bertrand SJ, Aksenova MV, Mactutus CF, Booze, RM (2013) HIV-1 Tat protein variants: Critical role for the cysteine region in synaptodendritic injury. Exp Neurol 248:228–235.

Bertrand SJ, Mactutus CF, Aksenova MV, Espensen-Sturges TD, Booze RM (2014) Synaptodendritic recovery following HIV Tat exposure: neurorestoration by phytoestrogens. J Neurochem 128:140–151

Bertrand SJ, Mactutus CF, Harrod SB, Moran LM, Booze, RM (2018) HIV-1 proteins dysregulate motivational processes and dopamine circuitry. Sci Rep doi:10.1038/s41598-018-25109-0

Bhatia MS, Munjal S. (2014). Prevalence of depression in people living with HIV/AIDS undergoing ART and factors associated with it. J Clin Diagn Res, 8:WC01–WC04.

Blanpied TA, Ehlers MD (2004). Microanatomy of dendritic spines: emerging principles of synaptic pathology in psychiatric and neurological disease. Biol Psychiatry 55:1121–1127

Booze RM, Wood ML, Welch MA, Berry S, Mactutus CF (1999) Estrous cyclicity and behavioral sensitization in female rats following repeated intravenous cocaine administration. Pharmacol, Biochem and Behav 64:605–610

Braestrup C, Sanchez C (2004). Escitalopram: a unique mechanism of action. Internat J Psy Clinical Practice:11–13.

Bryant VE, Whitehead NE, Burrell LE, Dotson VM, Cook RL, Malloy P, Devlin K., Cohen, R.A. (2015) Depression and apathy among people living with HIV: Implications for treatment of HIV associated neurocognitive disorders. AIDS Behav, 19:1430–1437

Budygin, E.A., Phillips, P.E.M., Robinson, D.L., Kennedy, A.P., Gainetdinov, R.R., & Wightman, R.M. (2001). Effect of acute ethanol on striatal dopamine neurotransmission in ambulatory rats. J Pharmacol Exp Thera, 297: 27–34.

Buyukdura JS, McClintock, SM Croarkin, PE (2013). Psychomotor retardation in depression: Biological underpinnings, measurement and treatment. Progress in Neuropsychopharm Bio Psyc, 35: 395–409

Campos LN, Guimaraes MD, Remien RH (2010). Anxiety and depression symptoms as risk factors for non-adherence to antiretroviral therapy in Brazil. AIDS Behav 14: 289–299.

Castellon SA, Hinkin CH, Wood S, Yarema KT (1998) Apathy, depression, and cognitive performance in HIV-1 infection. J Neuropsych 10:320–328

Chang L, Wang GJ, Volkow ND, Ernst T, Telang F, Logan J, Fowler JS (2008) Decreased brain dopamine transporters are related to cognitive deficits in HIV patients with or without cocaine abuse. Neuroimage 42:869–878

Denton AR (2019) Behavioral and Voltammetric Analysis of Chronic Escitalopram Treatment to the HIV-1 Transgenic Rat: Implications for Comorbid HIV-1 and Clinical Depression. (Master’s Thesis). Retrieved from https://scholarcommons.sc.edu/etd/5287

Denton AR, Samaranayake SA, Kirchner KN, Roscoe RF, Berger SN, Harrod SB, Mactutus CF, Hashemi P, Booze RM (2019) Selective monoaminergic and histaminergic circuit dysregulation following long-term HIV-1 protein exposure. J Neurovirology 25: 540–550

Di Matteo V, De Blasi A, Di Giulio C, Esposito E (2001). Role of 5-HT_2C_ receptors in the control of central dopamine function. Trends Pharmacol Sci 22:229–232.

Do AN, Rosenberg ES, Sullivan PS, Beer L, Strine TW, Schulden JD, Fagan JL, Freedman MS, Skarbinski J. (2014) Excess burden of depression among HIV-infected persons receiving medical care in the United States: Data from the medical monitoring project and the behavioral risk factor surveillance system. PloS One 9: e92842

Egenrieder L, Mitricheva E, Spanagel R, Noori HR (2020) No basal or drug-induced sex differences in striatal dopaminergic levels: a cluster and meta-analysis of rat microdialysis studies. J Neurochem 4:482–492

Ellis RJ, Calero P, Stockin MD (2009) HIV infection and the central nervous system: a primer. Neuropsychol Rev 19:144–151

Everall IP, Heaton RK, Marcotte TD, Ellis RJ, McCutchan JA, Atkinson JH, Grant I, Mallory M, Masliah E (1999) Corticalsynaptic density is reduced in mild to moderate human immunodeficiencyvirus neurocognitive disorder. Brain Pathol 9:209–217 (HNRC Group. HIV Neurobehavioral Research Center)

Farinpour R, Miller EN, Satz P, Selnes OA, Cohen BA, Becker JT (2003) Psychosocial risk factors of HIV morbidity and mortality: findings from the Multicenter AIDS Cohort Study (MACS). J Clin Exp Neuropsychol 25:654–670

Ferris MJ, Frederick-Duus D, Fadel J, Mactutus CF, Booze RM (2009) In vivo microdialysis in awake, freely moving rats demonstrates HIV-1 Tat-induced alterations in dopamine transmission. Synapse 63:181–185

Festa LK, Irollo E, Platt BJ, Tian Y, Floresco S, Meucci O (2020) CXXL12-induced rescue of cortical dendritic spines and cognitive flexibility. eLife 9:e49717.

Fitting S, Booze RM, Hasselrot U, Mactutus CF (2008) Differential long-term neurotoxicity of HIV-1 proteins in the rat hippocampal formation: a design-based stereological study. Hippocampus 18:135–147

Fitting S, Booze RM, Mactutus CF (2015) HIV-1 proteins, Tat and gp120, target the developing dopamine system. Curr HIV Res 13:21–42

Green MV, Raybuck JD, Zhang X, Wu MM, Thayer SA. (2019) Scaling synapses in the presence of HIV. Neurochem Res 44: 234–246.

Hashemi P, Dankoski EC, Petrovi J, Keithley RB, Wightman RM (2009) Voltammetric detection of 5-hydroxytryptamine release in the rat brain. Anal Chem 81:9462–9471

Henderson LJ, Johnson TP, Smith BR, Reoma LB, Santamaria UA, […] Nath A (2019) Presence of Tat and transactivation response element in spinal fluid despite antiretroviral therapy. AIDS 33:145–157

Horberg MA, Silverberg MJ, Hurley LB, Towner WJ, Klein DB, Bersoff-Matcha S (2008) Effects of depression and selective serotonin reuptake inhibitor use on adherence to highly active3 antiretroviral therapy and on clinical outcomes in HIV-infected patients. J Acquir Immune Defic Synr 47:384–390

Illenberger JM, Harrod SB, Mactutus, CF, McLaurin KA, Kallianpur, A, Booze RM (2020). HIV infection and neurocognitive disorders in the context of chronic drug abuse: Evidence for divergent findings dependent upon prior drug history. J Neuroimmune Pharm doi:10.1007/s11481-020-09928-5

Javadi-Paydar M, Roscoe RF, Denton AR, Mactutus CF, Booze RM (2017) HIV-1 and cocaine disrupt dopamine reuptake and medium spiny neurons in female rat striatum. PloSOne Doi:10.1371/journal.pone.0188404

Kesby JP, Markou A, Semenova S (2016) The effects of HIV-1 regulatory TAT protein expression on brain reward function, response to psychostimulants and delay-dependent memory in mice. Neuropharmacology 109:205–215

Kesby JP, Markou A, Semenova S (2016) Effects of HIV/TAT protein expression and chronic selegiline treatment on spatial memory, reversal learning and neurotransmitter levels in mice. Behav Brain Res 311:131–140.

Kesby JP, Najera JZ, Romoli B, Fang Y, Basova L, Birmingham A, Marcondes MCG, Dulcis D, Semenova S (2017) HIV-1 Tat protein enhances sensitization to methamphetamine by affecting dopaminergic function. Brain Behav Immun 65:210–221

Kim HJ, Martemyanov KA, Thayer SA (2008) Human immunodeficiency virus protein Tat induces synapse loss via a reversible process that is distinct from cell death. J Neurosci 28:12604–12613

Kim HJ, Shin AH, Thayer SA (2011) Activation of cannabinoidtype 2 receptors inhibits HIV-1 envelope glycoprotein gp120-induced synapse loss. Mol Pharmacol 80:357–366

Kumar AM, Ownby RL, Waldrop-Valverde D, Fernandez B, Kumar, M (2011) Human immunodeficiency virus infection in the CNS and decreased dopamine availability: relationship with neuropsychological performance. J Neurovirol 17:26–40

Marin RS, Firinciogullari S, Biedrzycki RC (1993) The sources of convergence between measures of apathy and depression. J Affect Disord 28:117–124

McIntosh RC, Rosselli M, Uddin L, Antoni, M (2015) Neuropathological sequelae of human immunodeficiency virus and apathy: A review of neuropsychological and neuroimaging studies. Neurosci Biobehav Rev 55:47–164

McLaurin KM, Booze RM, Mactutus CF (2016). Progression of temporal processing deficits in the HIV-1 transgenic rat. Sci Rep: doi:6:32831.10.1038/srep32831

McLaurin KM, Booze RM, Mactutus CF (2018) Evolution of the HIV-1 transgenic rat: utility in assessing the progression of HIV-1 associated neurocognitive disorders. J Neurovirol 24:229–245

McLaurin KA, Booze RM, Mactutus CF, Fairchild AJ (2017) Sex matters: Robust sex differences in signal detection in the HIV-1 transgenic rat. Front Behav Neurosci doi:103389/fnbeh.2017.00212

McLaurin KA, Cook AK, Li H, League AF, Mactutus CF, Booze RM (2018). Synaptic connectivity in medium spiny neurons of the nucleus accumbens: A sex-dependent mechanism underlying apathy in the HIV-1 transgenic rat. Front Behav Neurosci 12:285

McLaurin KA, Li H, Booze RM, Fairchild AJ, Mactutus CF (2018) Unraveling individual differences in the HIV-1 transgenic rat: Therapeutic efficacy of methylphenidate. Sci Rep 8:136

McLaurin KA, Li H, Booze RM, Mactutus CF (2019) Disruption of timing: NeuroHIV progression in the post-cART era. Sci Rep 9: 827

McLaurin KA, Mactutus CF, Booze RM, Fairchild AJ (2019) An empirical mediation analysis of mechanisms underlying HIV-1-associated neurocognitive disorders. Brain Res 1724:146436

McLaurin KA, Moran LM, Booze RM Mactutus (2020) Selective estrogen receptor beta agonists: a therapeutic approach for HIV-1 associated neurocognitive disorders. J Neuroimmune Pharmacol 15:264–279

Mills JC, Pence BW, Todd JV, Bengtson AM, Breger TL, Edmonds A (2018) Cumulative burden of depression and all-cause mortality in women living with HIV. Clin Infect Dis 67: 1575–1581

Moran LM, Booze RM, Mactutus CF (2013) Time and time again: Temporal processing demands implicate perceptual and gating deficits in the HIV-1 transgenic rat. J Neuroimmune Pharmacol 8: 988–997

Moran LM, McLaurin KA, Booze RM, Mactutus CF (2019) Neurorestoration of sustained attention in a model of HIV-1 Associated Neurocognitive Disorders. Front Behav Neurosci 13:169

Parsons LH, Justice JB (1993). Perfusate serotonin increases extracellular dopamine in the nucleus accumbens as measured by in vivo microdialysis. Brain Research 2:195–199

Paxinos G, Watson C (2014) The rat brain in stereotaxic coordinates. Academic Press, Elsevier

Pence BW, Mills JC, Bengtson AM, Gaynes BN, Berger TL, Cook RL (2018) Association of increased chronicity of depression with HIV appointment attendance, treatment failure, and mortality among HIV-infected adults in the United States. JAMA Psychiatry 74:379–85.

Purohit V, Rapaka R, Shurtleff D (2011) Drugs of abuse, dopamine, and HIV-associated neurocognitive disorders/HIV-associated dementia. Mol Neurobiol 44:102–110

Rabkin JG (2008). HIV and depression: 2008 review and update. Curr. HIV/AIDS Rep 5:163–171.

Reid W, Sadowska M, Denaro F, Rao S, Foulke J, Hayes N (…) Bryant J (2001) An HIV-1 transgenic rat that develops HIV-related pathology and immunologic dysfunction. Proc Natl Acad Sci U S A 98:9271–9276

Rodriguez A, Ehlenberger DB, Dickstein DL, Hof PR, Wearne SL (2008). Automated three-dimensional detection and shape classification of dendritic spines from fluorescene microscopy images. PlosOne 3, doi:10.1371/journal.pone.0001997

Roscoe R, Mactutus CF, Booze RM (2014. HIV-1 transgenic female rat: synaptodendritic alterations of medium spiny neurons in the nucleus accumbens. J Neuroimmune Pharmacol 9:642–653

Ruszczycki B, Szepesi Z, Wilczynski G, Bijata M, Kalita K, Kaczmarek L, Wlodarczyk J (2012). Sampling issues in quantitative analysis of dendritic spines morphology. BMC Bioinformatics 13:213

Sanmarti M, Ibanez L, Huertas S, Badenes D, Dalmau D, Slevein M, (…) Jaen A (2014) HIV-associated neurocognitive disorders. J Mol Psy, 2. Retrieved from http://www.jmolecularpsychiatry.com/content/2/1/2.

Savetsky JB, Sullivan LM, Clarke J, Stein MD, Sarnet JH (2001). Evolution of depressive symptoms in human immunodeficiency virus-infected patients entering primary care. J Nerv Ment Dis 189: 76–83

Saylor RA, Hersey M, West A, Buchanan AM, Berger SM, Nijhout HF, (…) Hashemi P. (2019) In vivo hippocampal serotonin dynamics in male and female mice: Determining effects of acute escitalopram using fast scan cyclic voltammetry. Front Neurosci 13: 362

Shaskan EG, Snyder SH (1970). Kinetics of serotonin accumulation into slices from rat brain: relationship to catecholamine uptake. J Pharmacol Exp Ther 175: 404–418

Sholl, DA (1953) Dendritic organization in the neurons of the visual and motor cortices of the cat. J Anat 87: 387–406

Silvers JM, Aksenov MY, Akensova MV, Beckly J, Olton P, Mactutus CF, Booze RM (2006) Dopaminergic marker proteins in the substantia nigra of human immunodeficiency virus type 1-infected brains. J Neurovirol 12:40–145

Sinharay S, Lee D, Shah S, Muthusamy S, Papadakis GZ, Ahang X, Maric D, Reid WC, Hammound, DA (2017) Cross-sectional and longitudinal small animal PET shows pre and post-synaptic striatal dopaminergic deficits in an animal model of HIV. J Nuc Med 55:27–33

Spudich S, Robertson KR, Bosch RJ, Gandhi RT, Cyktor JC, Mar H, […] Mellors JW (2019). Persistent HIV-infected cells in cerebrospinal fluid are associated with poorer neurocognitive performance. J Clin Invest 129:3339–3346

Strauss M, O’Donovan B, Ma Y, Xial Z, Lin S, Bardo MT, Ortinski PI, McLaughlin JP, Zhu J (2020) [3H} dopamine uptake through the dopamine and norepinephrine transporters is decreased in the prefrontal cortex of transgenic mice expressing HIV-1 transactivator or transcription. J Pharmacol Exp Ther 374:241–251

Toggas SM, Masliah E, Rockenstein EM, Rall GF, Abraham CR, Mucke L (1994) Central nervous system damage producedby expression of the HIV-1 coat protein gp120 in transgenicmice. Nature 367:188–193

Van Bockstaele EJ, Pickel VM (1993). Ultrastructure of serotonin-immunoreactive terminals in the core and shell of the rat nucleus accumbens: Cellular substrates for interactions with catecholamine afferents. J Comp Neurol 33:603–617.

Vigorito M, Connaghan KP, Chang SL (2015) The HIV-1 transgenic rat model of neuroHIV. Brain Behav Immun 48:336–349

West A, Best J, Abdalla A, Nijhout HF, Reed M, Hashemi P. (2018). Voltammetric evidence for discrete serotonin circuits, linked to specific reuptake domains, in the mouse medial prefrontal cortex. Neurochem Int 123:50–58.

Westwood FR (2008) The female rat reproductive cycle: a practical histological guide to staging. Toxicol Pathol 36:375–384

Yoo-Jeong M, Waldrop-Valverde D, Mccoy K, Ownby RL (2016) A structural equation model of HIV-related symptoms, depressive symptoms, and medication adherence. J HIV AIDS 2:1–15

Yuste, R (2010) Dendritic Spines. MIT Press

