## Supplementary figures and images for "Chronic SSRI treatment reverses HIV-1 protein-mediated synaptodendritic damage"

### Supplemental 1

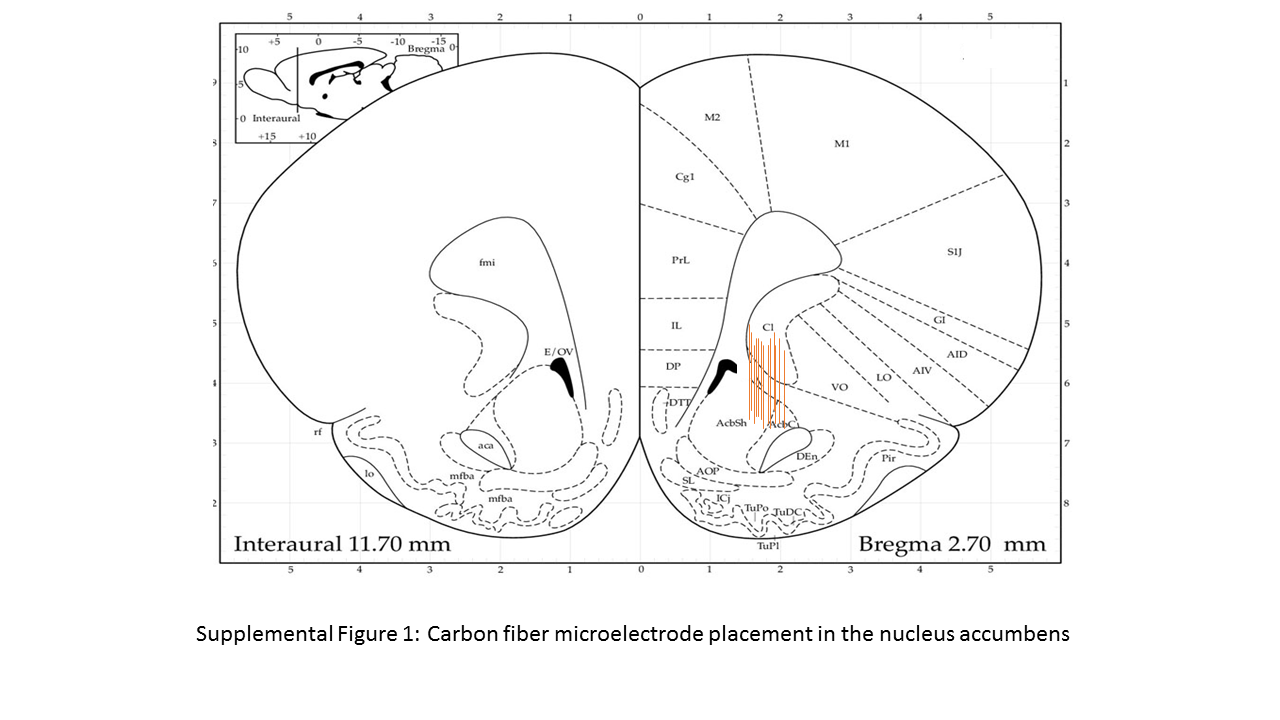

### Supplemental 2

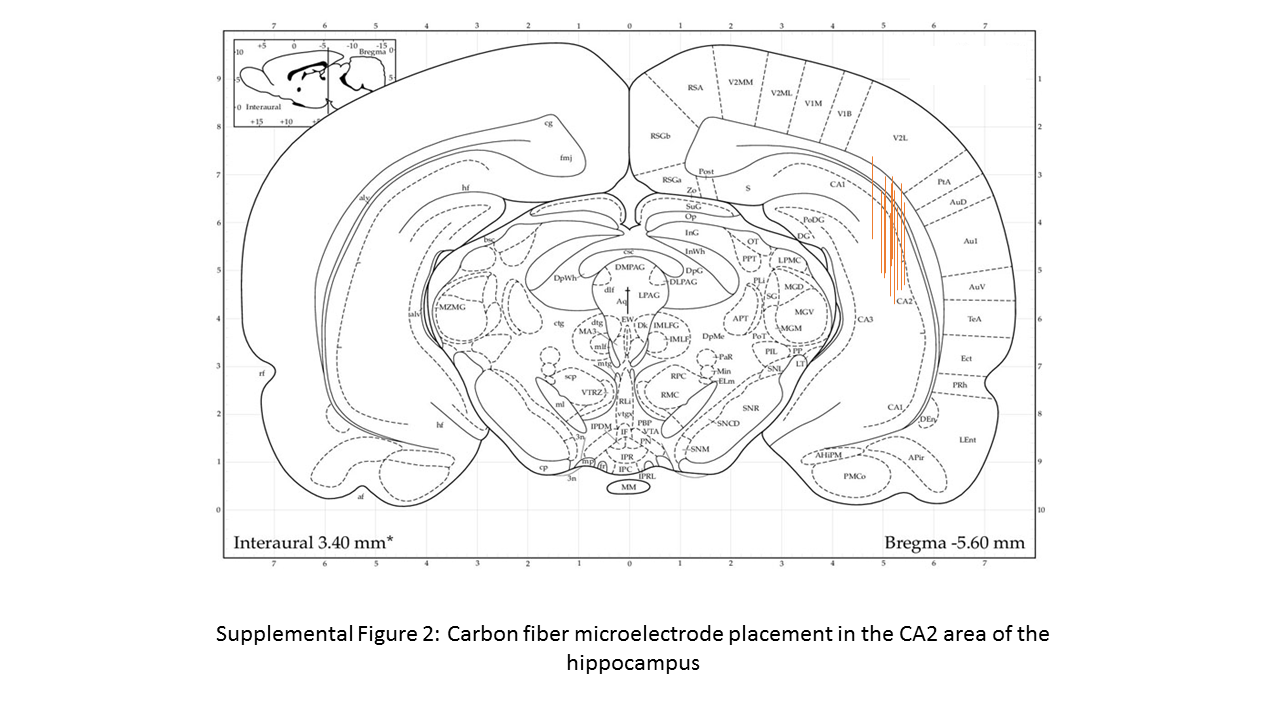

### Supplemental 3

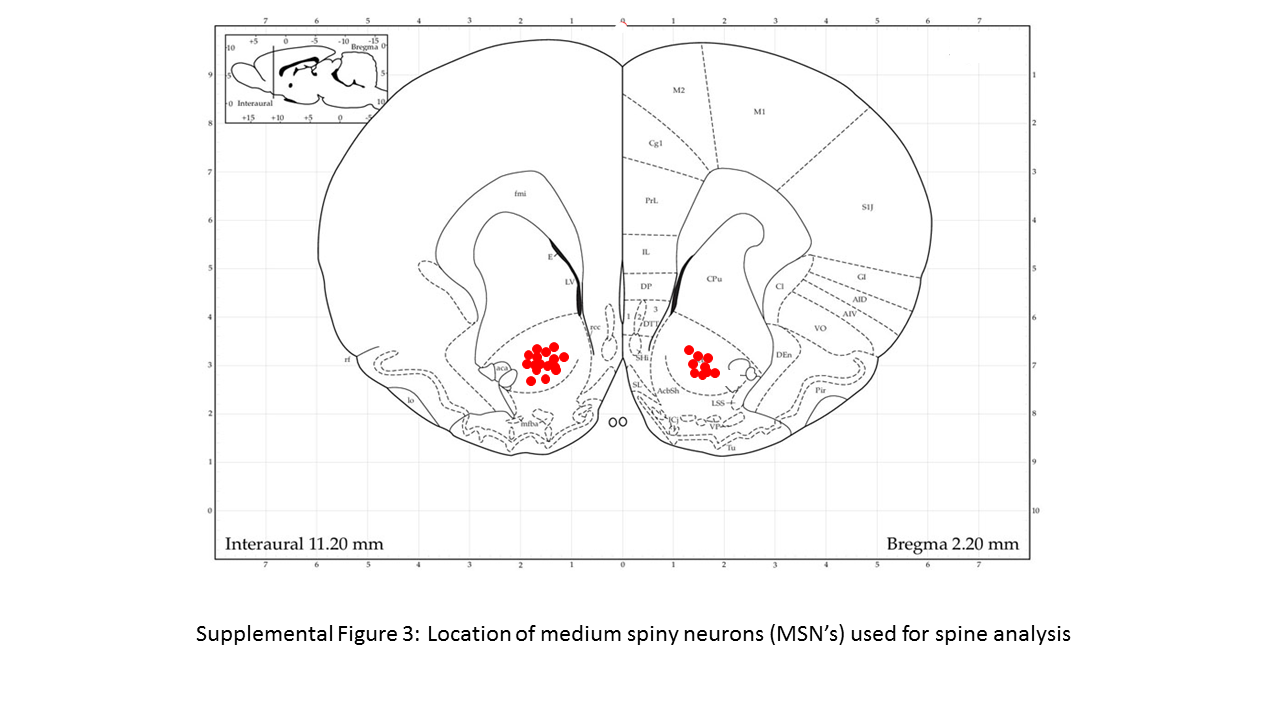
